# Maintaining their genetic distance; limited gene flow between widely hybridising species of *Geum* with contrasting mating systems

**DOI:** 10.1101/113969

**Authors:** Crispin Y. Jordan, Konrad Lohse, Frances Turner, Marian Thomson, Karim Gharbi, Richard A. Ennos

## Abstract

Mating system transition from outcrossing to selfing frequently gives rise to sister lineages with contrasting outcrossing rates. The evolutionary fate of such lineages depends on the extent to which they exchange genes. We measured gene flow between outcrossing *Geum rivale* and selfing *G. urbanum*, two sister species derived by mating system transition, which frequently hybridise. A draft genome was generated for *G. urbanum* and used to develop dd-RAD data scorable in both species. Coalescent analysis of RAD data from allopatric populations indicated that the two species diverged 2-3 Mya, and that long term gene flow between them has been very low (M=0.04). *G. rivale* showed greater genetic diversity in sympatry than allopatry, but genetic divergence between species was no lower in sympatry than allopatry, providing little evidence for recent introgression. Clustering of genotypes revealed that, apart from four early generation hybrids, individuals in sympatric populations fell into two genetically distinct groups with <1% admixture that corresponded exactly to their morphological species classification. Although our data suggest limited gene flow, we observed joint segregation of two putatively introgressed SNPs in *G. urbanum* populations that was associated with significant morphological variation; this provides tentative evidence for rare introduction of novel genetic diversity by interspecific gene flow. Our results indicate that despite frequent hybridisation, genetic exchange between *G. rivale* and *G. urbanum* has been very limited throughout their evolutionary history.

## Introduction

A key factor influencing the evolutionary trajectory of lineages is their level of gene exchange with related taxa (Abbott *et al.* 2016). A reduction in gene flow facilitates speciation (Coyne & Orr 2004). Limited gene flow from a related taxon may increase the genetic variability of a species and enable acquisition of novel traits that enhance its evolutionary potential, while maintaining its distinctness (Paoletti *et al.* 2006). However, if gene flow is more extensive, species may fuse (Carney *et al.* 2000). Thus, quantifying gene exchange among lineages is crucial for understanding the birth, continued evolution and possible extinction of species.

Quantifying gene flow between lineages is particularly valuable across lineage pairs that share an evolutionary transition, allowing patterns of genetic exchange associated with that evolutionary transition to be recognised. In many plant genera speciation is associated with a mating system transition from outcrossing to selfing (Stebbins 1957; Igic *et al.* 2006; Wright *et al.* 2013; Barrett *et al.* 2014). Although sister species with contrasting mating systems are typically ecologically distinct, hybridisation often occurs between them where habitat isolation breaks down or the ranges of the two taxa meet (e.g. *Mimulus* (Vickery, 1978; Brandvain *et al.* 2014; Kenny & Sweigart 2016), *Rhinanthus* (Ducarme *et al.* 2010), *Centaurium* (Brys *et al.* 2013)).

The transition from outcrossing to selfing affects genome evolution per se (Wright *et al.* 2008) and is commonly accompanied by changes in floral display (Sicard & Lenhard 2011). Thus the transition can potentially lead both to post-zygotic barriers to interspecific gene flow, due to interactions between differently evolved outcrossed and selfed genomes, and to pre-zygotic barriers, through its effect on mating opportunities (Fishman & Wyatt, 1999; Hu, 2015). Quantitative measurement of introgression between sister selfing and outcrossing taxa is needed to determine the strength of these effects, and to ascertain whether they prevent integration of novel genetic material into either species. Introgression would be particularly important for highly inbreeding taxa whose adaptive potential may be impaired by low genetic variability, and a reduced efficacy of positive and background selection (Charlesworth *etal.* 1993; Glémin & Ronfort 2013; Arunkumar *et al.* 2015).

Here we quantify gene flow between *Geum rivale* and *G. urbanum,* sister species that differ in mating system and associated floral display (Fig. 1). *Geum rivale,* which typically occupies open, moist habitats, is predominantly outcrossed (outcrossing rate t = 0.8) and possesses a pendulous flower adapted to bee pollination. In contrast, *G. urbanum,* a plant of shaded, well drained sites, is predominantly self-fertilised (t = 0.15) and bears erect flowers adapted to fly pollination (Taylor 1997a, b; Ruhsam *et al.* 2010).

**Figure 1.**
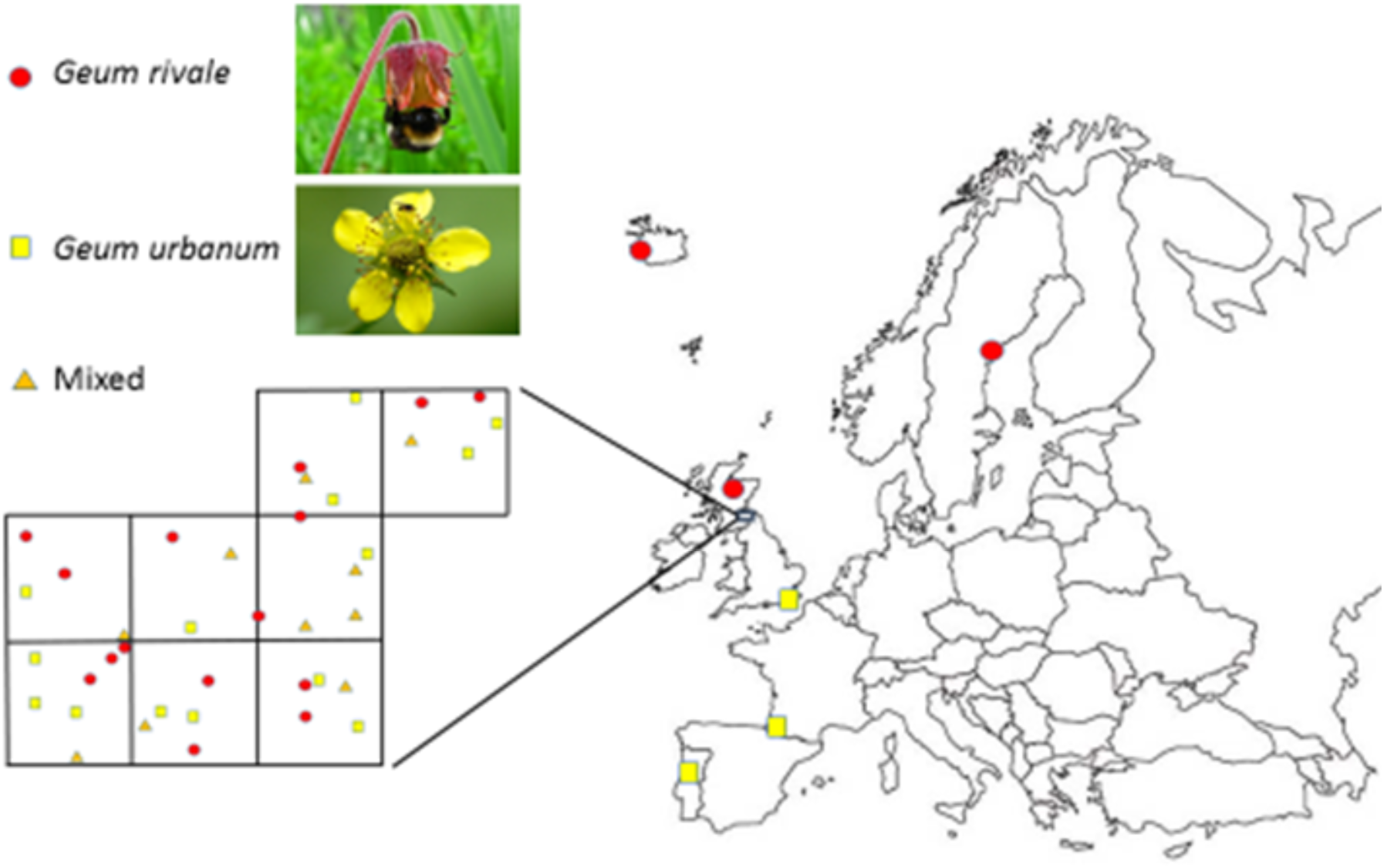
Locations of allopatric populations (large map) of *Geum rivale* and *G. urbanum* and detailed distribution of 39 sympatric populations in eight 10km x 10km squares (inset).

Allopatric populations of both species occur: *G. rivale* in tall herb montane communities in the UK, and at high latitude sites elsewhere in Europe; *G. urbanum* in the extreme south east of the UK, where rainfall is low, and at the southern extremes of its distribution in continental Europe (Taylor, 1997a, b). However, over most of their distributions in the UK and continental Europe the taxa are sympatric. Here they can be found as single species populations in “pure” habitats or together in “mixed” habitats. Typical mixed habitats include naturally wet woodlands and artificially disturbed river banks (Waldren *et al.* 1989).

In mixed habitats the highly fertile F1 hybrid *G. x intermedium* is widely reported, associated with hybrid swarms (Taylor 1997a). Detailed analysis of a single hybrid swarm indicated the presence of both F1 hybrids and early generation backcrosses to *G. rivale* that produce fertile seeds by both outcrossing and selfing (Ruhsam *et al.* 2011, 2013). The resulting recombinant offspring express no detectable fitness reduction when grown in a benign environment, but some recombinant types are not apparently present in the population of established adults in the hybrid swarm (Ruhsam 2013), perhaps due to selection (Ruhsam et al. 2013). Thus, the mating events occurring in mixed habitats are common and favourable for introgression, particularly via backcrossing to *G. rivale.* However, so far there has been no assessment of whether this has led to significant gene exchange between the two species.

To quantify gene exchange between the two *Geum* taxa we adopt a comparative genomic approach. We first generate a draft genome of the inbreeding species, *G. urbanum,* which we use to develop a set of SNP markers common to the two species. These SNPs are then scored in allopatric populations of both species likely to have been geographically isolated for the past 5ky. We analyse these data in a coalescent framework to estimate the age of the two taxa and measure interspecific gene flow from the time at which the species formed to the time when allopatric populations became isolated. We also use the allopatric samples to identify species-specific diagnostic SNPs.

To measure recent gene flow between the *Geum* taxa, in the period since the isolation of the allopatric populations, we analyse an additional set of samples taken from a broad area of sympatry. We look for signals of recent genome-wide introgression by comparing nucleotide diversity and genetic differentiation between the allopatric and sympatric samples. If introgression has occurred, significant increase in genetic diversity within and reduction in genetic differentiation between species are expected in sympatric compared to allopatric samples. We applied a clustering approach to the SNP data to determine the extent of admixture of *G. rivale* and *G. urbanum* genomes within individuals. Finally, we score the sympatric sample for previously identified species-specific diagnostic SNPs to quantify hybridisation and introgression at the level of individual genotypes. We relate these estimates of the hybridity of individuals, based on molecular data, to the morphology of these individuals measured in a common environment.

## MATERIALS AND METHODS

### Sampling

#### i. Sampling of Allopatric Populations

Four *G. rivale* individuals were sampled from each of three high elevation tall herb communities located on three distinct Scottish mountain ranges above the altitudinal limits of *G. urbanum* (Table S1; Taylor 1997b). One individual was also sampled from a single population in each of northern Sweden and Iceland where *G. urbanum* is absent (Table S1; http://linnaeus.nrm.se/flora).

Two *G. urbanum* plants were sampled from each of 10 populations in south-east England (Table S1), a region where *G. rivale* is absent due to lack of suitable habitats (Preston *et al.* 2002). Single individuals of *G. urbanum* were also sampled from two areas in Europe where *G. rivale* does not occur, namely Portugal and south-west France (Table S1; http://linnaeus.nrm.se/flora). 0.5 grms young leaf tissue was removed from sampled plants and stored dry in silica gel prior to DNA extraction.

#### ii. Sampling of Sympatric Populations

The Botanical Society of Britain and Ireland (BSBI) database was searched for the location of sites containing *G. rivale* alone (15), *G. urbanum* alone (14), or both taxa (10) in eight contiguous 10km x 10km squares within vice county 81 in the British Isles (Berwickshire) (Fig. 1, Table S2) (http://bsbidb.org.uk/maps/). At each site a single cutting was collected in August 2013 from each of two plants more than 5m apart, for each of the *Geum* taxa present. Field identification employed floral, fruit and vegetative characters. Newly emerging leaves (0.1 g.) were taken from each plant and frozen (−80°C) prior to DNA extraction. Cuttings were rooted in 20cm diameter pots containing commercial potting compost. Ninety six of the 98 cuttings were rooted successfully. Plants were overwintered outside in a randomised array in Edinburgh to induce flowering in spring 2014.

### Development of SNP markers via ddRAD

#### i. DNA extraction

DNA was extracted from dried samples using a modification of Doyle & Doyle (1990)’s CTAB protocol, while fresh and frozen material was extracted using the DNeasy plant mini prep kit (Qiagen) following the manufacturer’s protocol.

#### ii. Development of draft Genome

We developed a draft genome for the inbreeder, *G. urbanum* to facilitate SNP genotyping with double-digest Restriction Associated DNA sequencing markers (ddRAD; Peterson *et al.* 2012). We identified the individual within our Punnetts Town sample (Table S1) that was most homozygous at a set of 10 microsatellite loci (Arens *et al.* 2004). Using this sample we extracted 17μg of DNA from 150mg of young leaves. Details of the genome sequencing and assembly, which was conducted by Edinburgh Genomics, are given in the Supplementary Materials.

To identify SNPs within the draft genome and estimate coverage per scaffold, trimmed reads were mapped back to assembled scaffolds using bwa mem version 0.7.5a with -M option (Li & Durbin 2009). 94.0% of reads were aligned to scaffolds. Samtools (Li *et al.* 2009) version 0.1.18 was used to call variants.

#### iii. ddRAD library prep and sequencing

DNA quality was evaluated using the E-Gel Precast Agarose Electrophoresis System (ThermoFisher Scientific), and samples quantified using the Qubit dsDNA BR Assay Kit, (ThermoFisher Scientific). Sample preparation, library construction and PCR amplification for double digest RAD sequencing was modified from Peterson *et al.* (2012) (for full details see Supplementary Materials). Superpools of PCR products were sequenced either on an Illumina HiSeq 2500 using 125 based paired-end reads in high output mode (v4 chemistry) or an Illumina MiSeq using 150 base paired-end reads (v2 chemistry).

#### iv. Quality filtering and preparation of reads from ddRAD

The bioinformatics and analysis pipeline for all ddRAD data is summarized in Figure S1. To match the read lengths produced by MiSeq and HiSeq Illumina technologies for ddRAD analyses, we used fastx trimmer (http://hannonlab.cshl.edu/fastx_toolkit/) to trim MiSeq reads to 125bp (STACKS requires reads of equal length). We then de-multiplexed and filtered reads for quality using process_radtags (STACKS v 1.21; Catchen *et al.* 2011, 2013). This process removed reads with an uncalled base, and those with an average quality score below 20 over a sliding window comprising 15% of a read. Finally, we again trimmed reverse reads to 117bp with fastX trimmer to produce forward and reverse reads of equivalent length after the 8bp barcode was removed from the forward reads.

#### v. Aligning reads to draft genome

Both *Geum* species are ancient hexaploids (2N = 42), with disomic inheritance (Smedmark *et al.* 2003; Vandepitte *et al.* 2007; Ruhsam *et al.* 2010). Mapping ddRAD sequences from the two *Geum* species must account for divergence between species to allow reads from both species to map to the *G. urbanum* draft genome. However, allowing an excessive number of mismatches can cause reads to map to multiple loci in ancient polyploid taxa, like *Geum.* We explored the effect of varying the number of allowed mismatches (M) on alignments for one allopatric individual from each species (Punnetts Town, *G. urbanum;* Ben Lui, *G. rivale;* both sequenced with Illumina HiSeq).

For a range of M values, we used GSnap (version 2014-12-16; Wu & Watanabe 2005) to map reads to the draft genome. We required that reads map uniquely, specified an indel penalty of two, and did not allow terminal alignments. For the *G. urbanum* sample, between 64.86% to 67.74% of paired-end (PE) ddRAD reads mapped as proper pairs when we allowed one to nine mismatches (M=1 to 9); however the percentage of reads that aligned jumped to 82.84% for M=10. Similarly, the percentage of *G. rivale* PE reads that aligned in proper pairs increased gradually from 39.21% to 58.16% over M=2 to M=9, with a large increase to 79.93% for M=10. Therefore, the structure of the *Geum* genome (e.g., due to polyploidy and/or repetitive sequences) appears to present different environments for aligning ddRAD data at a threshold genetic diversity of ~8.55% (i.e., 10/117 * 100, where 117 equals the read length). Consequently, we performed all subsequent alignments as above, setting M=6 to avoid this threshold but also allow many reads from *G. rivale* to map to the heterospecific draft genome.

#### vi. Assembling radtags and calling SNPs

We used STACKS’ ref_map pipeline (v 1.21; Catchen *et al.* 2011, 2013; Hohenlohe *et al.* 2010) to assemble the aligned reads into radtags. Next, we used STACKS’ rxstacks module to correct genotype calls based on population-level genotype information using a bounded (error rate) model with the parameters conf_lim and bound_high equal to 0.25 and 0.1, respectively (see http://catchenlab.life.illinois.edu/stacks/manual/).

#### vii. Identifying and filtering for paralogs

Ancient hexaploidy complicates analyses of *Geum* because reads from paralogous genome regions could map to identical sites. As a strict filter for identifying genome regions that potentially attract paralogous reads, we used STACKS’ populations module to calculate observed heterozygosity and F_IS_ at each identified SNP. Thereafter custom scripts were used to collate data and restrict further analysis to scaffolds free of SNPs that exhibited either excess heterozygosity (>0.5) or negative F_IS_.

The heterozygosity and F_IS_ calculations *for this purpose only* considered UK allopatric *Geum* samples that comprised four individuals from each of three Scottish ‘allopatric’ *G. rivale* populations (12 *G. rivale* individuals), and one randomly chosen individual from each of the 10 Southern England *G. urbanum* populations (Table S1). Nucleotide-level heterozygosity and F_IS_ were calculated separately for *G. rivale* and *G. urbanum;* we assumed each set of samples was from a single, large UK population, despite population structure being evident in both species (see Results). Population structure reduces the number of heterozygotes compared to a single panmictic population: therefore, our filters should preferentially remove genomic regions with excess heterozygosity due to paralogy.

### Population Genetic Analysis of Allopatric Populations

#### i. Patterns of Polymorphism within and between species

We characterized polymorphism within and between the *Geum* species using the 12 allopatric UK *G. rivale* samples plus one from each of Iceland and Sweden, and 10 UK *G. urbanum* samples (one from each allopatric UK population) plus one each from France and Portugal (Table S1). We used STACKS’ populations module to identify SNPs that are polymorphic within at least one species or show a fixed difference between species. This analysis only considered radtags present in all 26 individuals described above.

#### ii. Inbreeding coefficients and population differentiation

We applied the STACKS’ populations module to estimate the inbreeding coefficients within each population (F_IS_) and measure differentiation among populations (F_ST_) for each species using data from the three allopatric *G. rivale* populations in Scotland (4 individuals/pop) and the ten allopatric *G. urbanum* populations in England (2 individuals/pop). Estimates of F_IS_ and F_ST_ only considered radtags present in both species, and that were present in all individuals of a given analysis.

#### iii. Identification of species-specific SNPs

We used alternately fixed SNPs in allopatric populations of the two taxa (identified in (i.) above) to provide an initial list of species-specific SNPs. To minimize linkage between diagnostic SNPs and obtain an estimate of introgression across the whole genome, we selected one (the first) alternately fixed SNP per scaffold for the introgression analysis. However, we note that by using the first SNP per scaffold, we bias our data against (larger) well-assembled scaffolds. Therefore, we repeated our analyses of introgression using all available species-diagnostics SNPs to test for an effect of such bias, and obtained qualitatively identical results (not shown).

#### iv. Coalescent analysis of gene flow during lineage divergence

We used an analytic likelihood framework to assess the support for alternative models of divergence between *G. rivale* and *G. urbanum* with and without gene flow. The method is described in Lohse *et al.* (2016). Briefly, the analysis is based on a single diploid individual for each species and considers the blockwise site frequency spectrum, i.e. the joint frequencies of four polymorphism types (as in “*Patterns of Polymorphism within and between species*”, above) in short blocks of sequence: i) heterozygous sites exclusive to *G. rivale*, ii) heterozygous sites exclusive to *G. urbanum*, iii) heterozygous sites shared by both species and iv) fixed differences between species. We counted these site types within 117bp radtags (block), and treated each radtag as an independent block. For randomly mating populations, the polymorphisms at each block represent an independent outcome of the coalescent process, which is a function of the species’ history. Since STACKs ignores RAD tags that are monomorphic, we conditioned the likelihood on only observing variable blocks. To do this, we normalized the probability of each blockwise mutational configuration by 1-p_IBS, where p_IBS is the probability of identity in state for a block.

We primarily wish to test for historical introgression between *G. rivale* and *G. urbanum.* Therefore, we compared three models: (1; “div_2_”) species diverged at time *T* with no introgression, (2; “IM_u→r_”) a history of divergence with gene flow from *G. urbanum* to *G. rivale* and (3; “IM_r→u_”) divergence with gene flow from *G. rivale* to *G. urbanum* (4 parameters). All models assume instantaneous species divergence and a constant N_e_ within taxa. We also constrained the effective size of the ancestor to equal that of *G. rivale*. To capture the decrease of N_e_ expected to result from the transition to inbreeding (Charlesworth & Wright 2001), we allowed N_e_ to differ between *G. urbanum* and the other taxa (the ancestor of *G. rivale* and *G. urbanum* is most likely outcrossing given the rarity of transitions from self-compatibility to self-incompatibility; Igic et al. 2006). Migration was modelled as a constant rate M = 4N_anc_ x m individuals per generation. We converted estimates of T into years using t = T × 2N × g where g is generation time and N = θ/4μ. We assumed a mutation rate estimated for *Arabidopsis thaliana*, μ = 7 × 10^-9^ (Ossowski *et al.* 2010), and an average generation time across the two species of 3 years (Taylor 1997a, b) (generations are likely longer in *G. rivale* than in *G. urbanum*).

Inbreeding complicates this analysis because it reduces genetic diversity within individuals relative to the population level. To minimize the confounding effect of inbreeding on the analyses, we initially chose the most outbred individuals for analysis. We used a recently outcrossed *G. urbanum* individual (see Results; population Mill Wood), which had genetic diversity (π = 0.0009 ± 0.0001) similar to the UK allopatric *G. urbanum* populations as a whole (see Results) and so may adequately represent the diversity of this species in the UK. “Leaky” self-incompatibility also introduces variation in inbreeding among individuals of *G. rivale.* Therefore, we conducted separate analyses that paired the outbred *G. urbanum* Mill Wood individual with 3 allopatric UK samples of *G. rivale* that span a range of heterozygosity to examine how inbreeding in *G. rivale* affects our conclusions. Maximum likelihood estimates under each model were obtained in *Mathematica* v. 10.2 (see File S2 of Lohse *et al.* 2016). To estimate 95% CI for M and T, we obtained discretized marginal support (logarithm of the likelihood) curves for these parameters (maximising the likelihood for all other parameters at each point).

### Population Genetic Comparison of Allopatric and Sympatric Populations

#### i. Genetic diversity in allopatric and sympatric populations

To determine whether potential genetic exchange in sympatric populations resulted in an increase in genetic diversity, we estimated genetic diversity (π) in UK allopatric and Berwickshire samples (pooling Berwickshire’s ‘pure’ and ‘mixed’ populations) of each species using STACKS’ populations module (providing mean and standard error, S.E.). The analysis considered radtags present in at least half of individuals for each species/population type (allo-vs sympatric) combination; we obtained qualitatively similar results when we restricted the analysis to radtags present in all individuals (not shown). The analysis excluded four individuals in the Berwickshire sample identified as obvious early generation hybrids or backcrosses (see Results). This analysis therefore involved 12 allopatric *G. rivale*, 20 allopatric *G. urbanum*, 48 sympatric *G. rivale* and 45 sympatric *G. urbanum* samples.

#### ii. Genetic differentiation between taxa in allopatry and sympatry

To test whether potential gene exchange resulted in less differentiation between the species in sympatric populations, we calculated genetic differentiation d_xy_ = ∑[(PX * (1−PY)) + ((1−PX) * PY)] / n between the two species, separately for allopatric and sympatric samples: P represents the frequency of a focal allele in the sample of species X and Y, and n is the sequence length (Nei & Li 1979; Cruickshank & Hahn 2014). These calculations considered the same samples as used to calculate genetic diversity (π), described above. We estimated d_xy_ for each scaffold, which we assume to be independent with respect to linkage and determined the mean d_xy_ and its S.E. across scaffolds. All analyzed radtags were present in at least 12 individuals for each species/population type combination, and the same scaffolds were analyzed in allopatric and sympatric estimates of d_xy_ (n=418 scaffolds).

### Genetic Analysis of Hybrids and Introgression in Sympatric Populations

#### i. Cluster analysis using fastSTRUCTURE

As a first approach to analyzing introgression between the two *Geum* taxa in sympatry, we used genotypic clustering implemented in fastSTRUCTURE (Raj *et al.* 2014). Our analysis assumed that samples derived from two populations (i.e., K=2) corresponding to the two *Geum* species, with the possibility of genetic admixture of individuals. The SNP data was derived from a STACKS analysis of all 92 successfully genotyped individuals in the sympatric Berwickshire sample together with data from the British and European allopatric populations of the two species. We analyzed two datasets that differed in degree of filtering: (1) SNPs were present in any fraction of individuals, which led to greater numbers of SNPs but missing values for some individuals, and (2) all SNPs were present in all individuals. Both analyses used SNPs that had been filtered for paralogs, and considered only a single SNP per scaffold to minimize the effect of linkage between SNPs.

#### ii. Identifying hybrids and introgressed individuals using species-specific SNPs

In our second approach to analyzing recent introgression we used custom scripts and the species-specific SNPs identified in (iii.) (above) to estimate the fraction of alleles in each individual within the sympatric Berwickshire sample that is *G. rivale* in origin (Hybrid Index). SNPs with either un-callable genotypes (based on STACKS’ likelihood algorithm) or third alleles (e.g., due to sequencing or alignment error) were excluded from the calculations. We simultaneously tabulated the frequency of SNPs that were heterozygous for the species-diagnostic alleles in each individual.

In general, our analyses did not specify minimum coverage because STACKS accounts for coverage when calling genotypes (or leaving a genotype uncalled) (Hohenlohe *et al.* 2010; Catchen *et al.* 2013). However, to check whether specifying minimum coverage would alter our results, we repeated our analyses, requiring a minimum coverage of 20 in all analysed individuals when running fastSTRUCTURE and creating the panel of species-diagnostic SNPs. This quartered the number of species-diagnostic SNPs, but yielded qualitatively identical results (not shown).

### Analysis of Morphological Variation in Sympatric Populations

#### i. Measurement of morphological variation in sympatric populations

From the beginning of April 2014 plants grown from cuttings from sympatric populations were monitored weekly for flowering. From each plant a newly opened flower and the stipule located on the flowering stem immediately below the flower were sampled. The following characters, which discriminate between *G. rivale* and *G. urbanum*, were measured: angle at which flowers are held (degrees from vertical) (FA), petal length (PL) (mm), petal width (PW) (mm), petal shape (PS) (proportional height of widest part of petal)), sepal length (SL) (mm), stamen number (SN), stipule length (STL) (mm) and stipule width (STW) (mm) (Ruhsam *et al.* 2011, 2013).

#### ii. Statistical analysis of morphological data

Principal Component Analysis (PCA) was conducted on the total sympatric sample using data on all eight morphological characters measured. Morphologically distinct groupings were recognised on scatterplots of the first two principal component scores and related to parental species and hybrid classes defined genetically by species-specific SNPs. PCA based on the same characters was also used to summarise morphological variation separately within *G. urbanum*. GLMs were used to test for the effect of two SNP variants, putatively jointly introgressed from *G. rivale* (see results), on the first two principal component scores for *G. urbanum.* All statistical analyses were conducted in Minitab 16.

## RESULTS

### Development of SNP markers via ddRAD

#### i. Draft Genome

The genome assembly contained a total of 170,030 scaffolds, with an N50 of 24.6Kb and total assembly size of 1.2Gb. In order to identify signs that scaffolds represent multiple copies of the genome, the distributions of coverage per scaffold and percent variant bases per scaffold was assessed. Scaffolds of length less than 10Kb were excluded, leaving 32,182 scaffolds (with a total length of 909 Mb). The distribution of percent variant bases per scaffold is shown in Figure S2, and the distribution of coverage per scaffold is shown in Figure S3. Coverage per scaffold follows a roughly normal distribution, which would be expected if the scaffolds mostly represented the same number of copies of the genome.

Core genic regions were well assembled, and appeared to be present in approximately three (haploid) copies as expected for an ancient hexaploid. We searched our assembled genome for 248 ultra-conserved eukaryotic genes (CEGs), listed by Parra *et al.* (2007), using their Core Eukaryotic Genes Mapping Approach. We identified 93% and 97% of the core genes that were complete or partially complete, respectively. On average, complete and partially complete CEGs were represented by 3.39 and 3.82 orthologs per CEG, respectively, with at least 90% of CEGs represented by more than one ortholog.

#### ii. ddRAD tags

Following quality filtering, 2.7 * 10^7^ reads remained in the MiSeq data, and between 6.0 * 10^7^ and 7.3 * 10^7^ reads remained that derived from the five HiSeq libraries. Following alignments, assembling radtags with STACKS, and applying corrections with rxstacks, our dataset included 230,356 radtags for *G. rivale* and *G. urbanum,* collectively. However, coverage was highly stochastic. For example, only ~2% (4524) of radtags were represented in more than one half of our samples.

#### iii. Identifying and filtering for paralogs

When the STACKS’ populations module was used to analyse the raw SNP data from the allopatric populations of the inbreeding taxon *G. urbanum*, F_IS_ estimates were low and sometimes negative, which is unexpected for a highly inbreeding species. This suggests that paralogous reads have mapped to identical locations and thereby increased individual heterozygosity (see Table S4). We therefore applied our filter for paralogy (rejecting 1344 scaffolds with SNPs that exhibit either excess heterozygosity (>0.5) or negative F_IS_) and, unless specifically noted, we henceforth only present results from paralogy-filtered SNPs.

### Population Genetic Analysis of Allopatric Populations

#### i. Patterns of polymorphism within and between species

The majority of SNPs in the dataset were polymorphic in *G. rivale* but invariant in *G. urbanum.* This type of polymorphic site occurred approximately four times more frequently than the reverse case (Table 1). 22% of SNPs were alternately fixed between the species (Table 1) and only 1.5% of SNPs were shared polymorphism (Table 1).

**Table 1:** Frequency of polymorphism of different forms within and between *Geum* species. Analysis allows multiple SNPs per scaffold.

| Polymorphism Type | Count |
| --- | --- |
| Alternately fixed SNPs | 488 |
| SNP polymorphic in *G. rivale* but one allele fixed in *G. urbanum* (e.g., *G. rivale*: A, T; *G. urbanum*: T) | 1334 |
| SNP polymorphic in *G. urbanum* but one allele fixed in *G. rivale* | 338 |
| *G. rivale* and *G. urbanum* share polymorphism | 34 |
| Total SNPs analysed | 2194 |

#### ii. Inbreeding coefficients and population differentiation

For the allopatric populations sampled in Britain we obtained F_IS_ estimates close to 0.25 for *G. rivale,* consistent with leaky self-incompatibility (Ruhsam *et al.* 2010), and F_IS_ greater than 0.9, consistent with very high selfing rates, for eight of the ten UK ‘pure’ *G. urbanum* populations (Table 2). Among the two *G. urbanum* populations with low F_IS_, the first population (Mill Wood F_IS_ = 0.0845), included one (of the two) sample(s) that was heterozygous at 92% of the 71 polymorphic SNPs analysed, suggesting that this sample was derived from a recent outcrossing event between two diverged inbred lines. In the second population, Selwyn Wood, only one SNP was recorded as polymorphic, suggesting that this population may have been founded by few (possibly a single) highly selfing individuals.

**Table 2.**
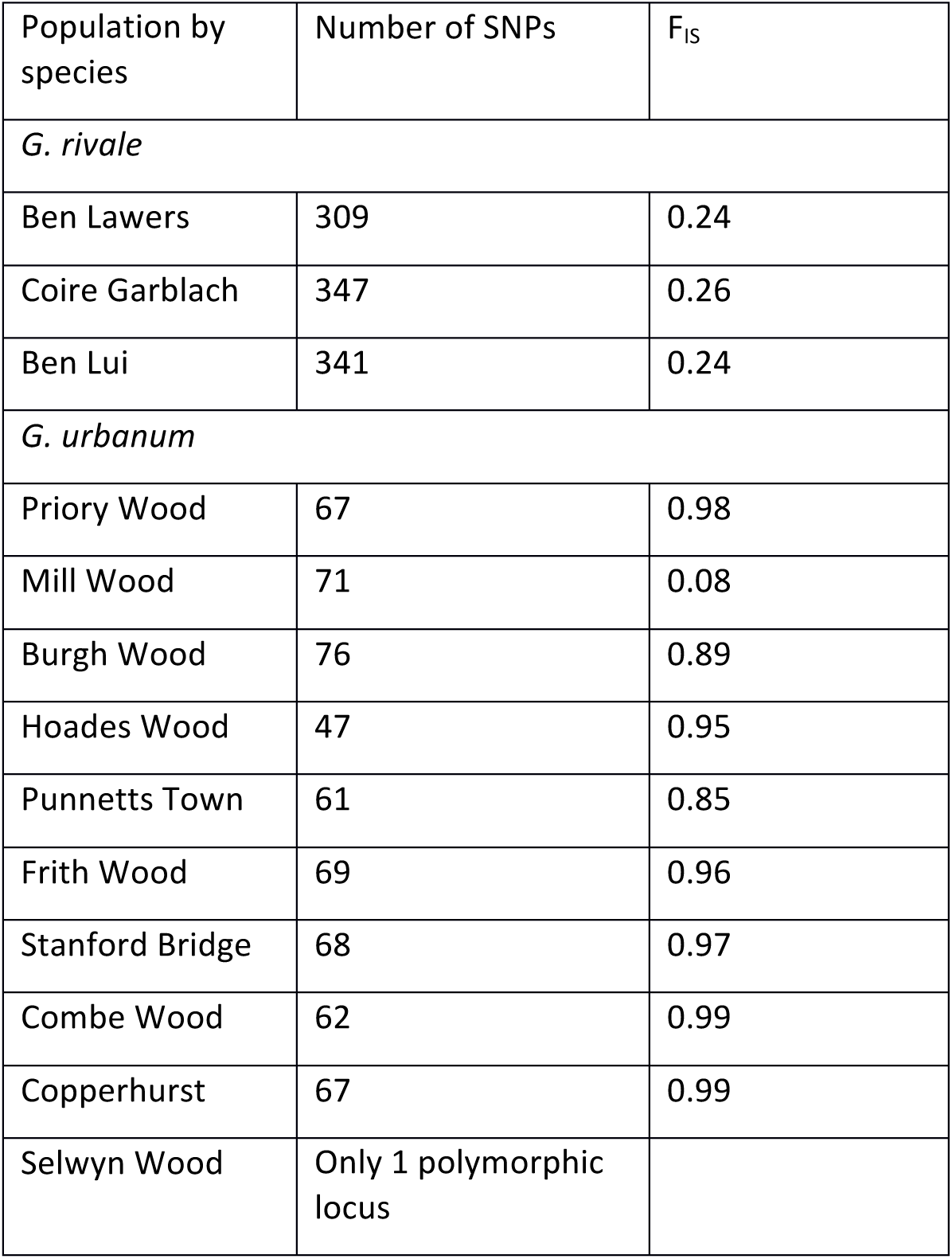
F_IS_ estimates for ‘allopatric’ UK *G. rivale* and *G. urbanum* populations using data filtered for paralogs.

Allopatric populations of *G. rivale* in Britain exhibited less population differentiation than *G. urbanum* populations (mean pairwise F_ST_ 0.13 and 0.38 respectively), as expected from their contrasting breeding systems.

#### iii. Coalescent analysis of gene flow during lineage divergence

Table S5 summarizes the numbers of each polymorphism type and blocks analysed for the *G. urbanum* - *G.rivale* sample pairs. All three pairs had approximately 1400 SNPs distributed among ca. 660 blocks (i.e., radtags). Model comparisons for all three sample pairs reject a model of strict divergence, and suggest that introgression has occurred between the *Geum* species (Table 3), but at a very low rate (see below). For two sample pairs (involving *G. rivale* samples Ben Lui 1 and Ben Lawers 5), the model of gene flow from *G. rivale* to *G. urbanum* (i.e., IM_r→u_) fit the data significantly better than the model of strict divergence (i.e., div_2_), whereas the model IM_u→r_ does not fit significantly better than div_2_ (Table 3). Models that include gene flow (IM_r→u_ and IM_u→r_) also fit the data significantly better than the div_2_ model for the third pair (involving sample Ben Lui 4), but IM_r→u_ and IM_u→r_ have effectively equal support (Table 3). Results from this latter pair likely differ from the former pairs because it includes a single block with a shared heterozygous site, while the other sample pairs lack shared heterozygous sites (Tables S5, S6). As Ben Lui 4 is likely the most heterozygous *G. rivale* sample (Table S5), we focus on this pair, but note that parameter estimates (Table S6) and general conclusions (i.e., support for very low introgression rates) are similar for all three sample pairs.

**Table 3.** Difference in log Likelihoods between the “best” and alternate models. “Sample” refers to which *G. rivale* individual is paired with the single *G. urbanum* individual

| Sample | $\text{div}_2$ | $\text{IM}_{u \rightarrow r}$ | $\text{IM}_{r \rightarrow u}$ |
| --- | --- | --- | --- |
| Ben Lui 1 | -12.9 | -12.5 | 0 |
| Ben Lui 4 | -9.66 | 0 | -0.721 |
| Ben Lawers 5 | -6.67 | -6.51 | 0 |

The three models (div_2_, IM_r→u_, IM_u→v_) yield similar estimates of Ne and divergence time (Ben Lui 4 pair: Table 4; all three sample pairs: Table S6). In general, Ne of *G. urbanum* is a half to a quarter of that of *G. rivale* (and their common ancestor), and all three models suggest that the species diverged approximately 2 to 3 million years ago (Table 4; Table S6). Models IM_u→v_ and IM_r→u_ both suggest a low but significant long-term rate of effective gene flow (M ≈ 0.04), of approximately one migrant every 25 generations (Table 4; see Table S6 for M for additional sample pairs).

**Table 4.**
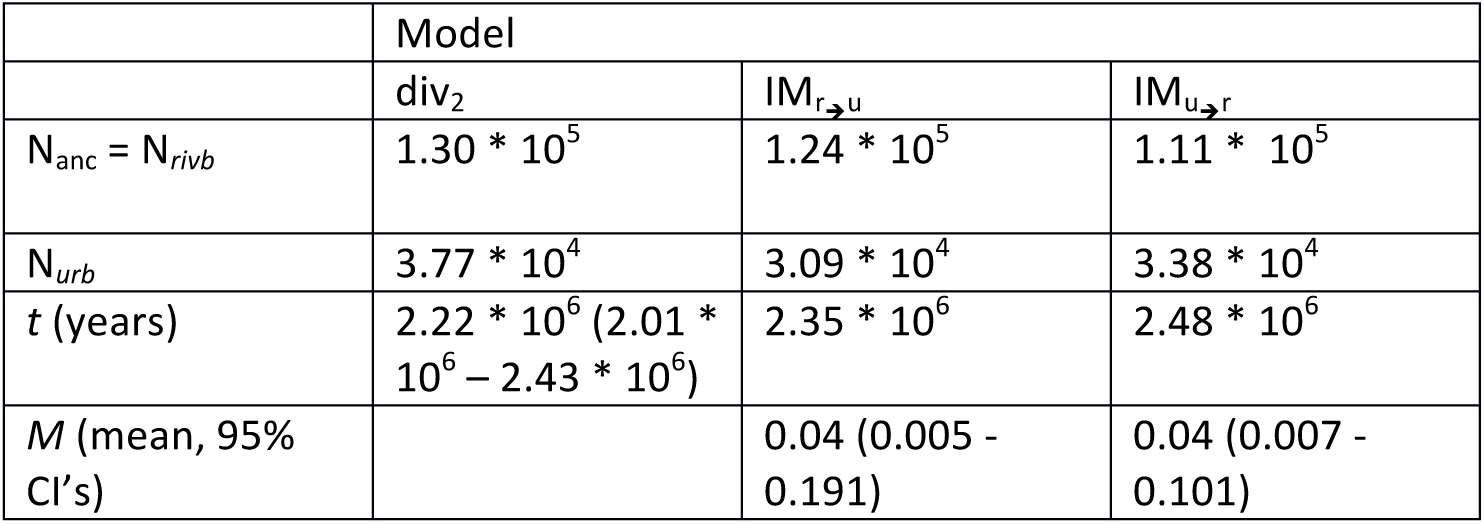
Parameter estimates under the models for *G. rivale* sample, Ben Lui 4. 95% Confidence intervals provided in parentheses. See Table S6 for parameter estimates for all three sample pairs.

|  | Model |  |  |
| --- | --- | --- | --- |
| | $\text{div}_2$ | $\text{IM}_{r \rightarrow u}$ | $\text{IM}_{u \rightarrow r}$ |
| $N_{\text{anc}} = N_{\text{rivb}}$ | $1.30 * 10^5$ | $1.24 * 10^5$ | $1.11 * 10^5$ |
| $N_{\text{urb}}$ | $3.77 * 10^4$ | $3.09 * 10^4$ | $3.38 * 10^4$ |
| $t$ (years) | $2.22 * 10^6$ ($2.01 * 10^6 - 2.43 * 10^6$) | $2.35 * 10^6$ | $2.48 * 10^6$ |
| $M$ (mean, 95% CI's) | | 0.04 (0.005 - 0.191) | 0.04 (0.007 - 0.101) |

### Population Genetic Comparison of Allopatric and Sympatric Populations

#### i. Genetic diversity in allopatric and sympatric populations

Estimates of genetic diversity (π) were based on 826 radtags, each 117 base pairs in length, totaling 96 642 base-pairs; as these radtags were the same as those used to develop species-diagnostic SNPs, they occur in both species. Genetic diversity was greater in *G. rivale* (mean π ± SE: ‘allopatric’ 0.0044 ± 0.0001; Berwickshire 0.0048 ± 0.0001) than *G. urbanum* (‘allopatric’ 0.0011 ± 0.0001; Berwickshire 0.0011 ± 0.0001), and was similar between population types for *G. urbanum*. However *G. rivale* genetic diversity was slightly greater for sympatric than allopatric samples. Since STACKS ignores monomorphic radtags, we note that these estimates of π should be interpreted in terms of relative, but not absolute diversity (see also Arnold *et al.* 2013).

#### ii. Genetic differentiation between taxa in allopatry and sympatry

Genetic differentiation between *G. rivale* and *G. urbanum* is similar for UK allopatric and sympatric samples (mean d_xy_ (+/- SE) equals 0.0115 +/- 0.0005 and 0.0112 +/- 0.0005, respectively). Again, we note that, due to the fact that STACKS’ ignores monomorphic radtags, these estimates should be viewed as relative measures (and upper limits) of d_xy_, and not as absolute estimates.

### Hybrids and Introgression in Sympatric Populations

#### i. Cluster analysis using fastSTRUCTURE

*fastSTRUCTURE* analysis that considered SNPs present in either all (188 SNPs) or a proportion of (492 SNPs; results not shown) individuals analysed in the combined allopatric and sympatric populations yielded highly consistent results. Figure 2 illustrates fastSTRUCTURE results for analyses based on 188 SNPs. All except one of the 36 individuals in the allopatric populations, and all except 4 of the 96 individuals in the sympatric populations, show less than 1% admixture. In the sympatric population Q values (proportion of *G. rivale* genome) of the four individuals with substantial admixture are 0.496, 0.517, 0.663 and 0.935. On the basis of these Q values there is no evidence for significant introgression from *G. rivale* into *G. urbanum.* The substantially admixed individuals present are likely to be recently formed F1 or F2 hybrids, and different generations of backcrosses to *G. rivale.*

**Figure 2.**
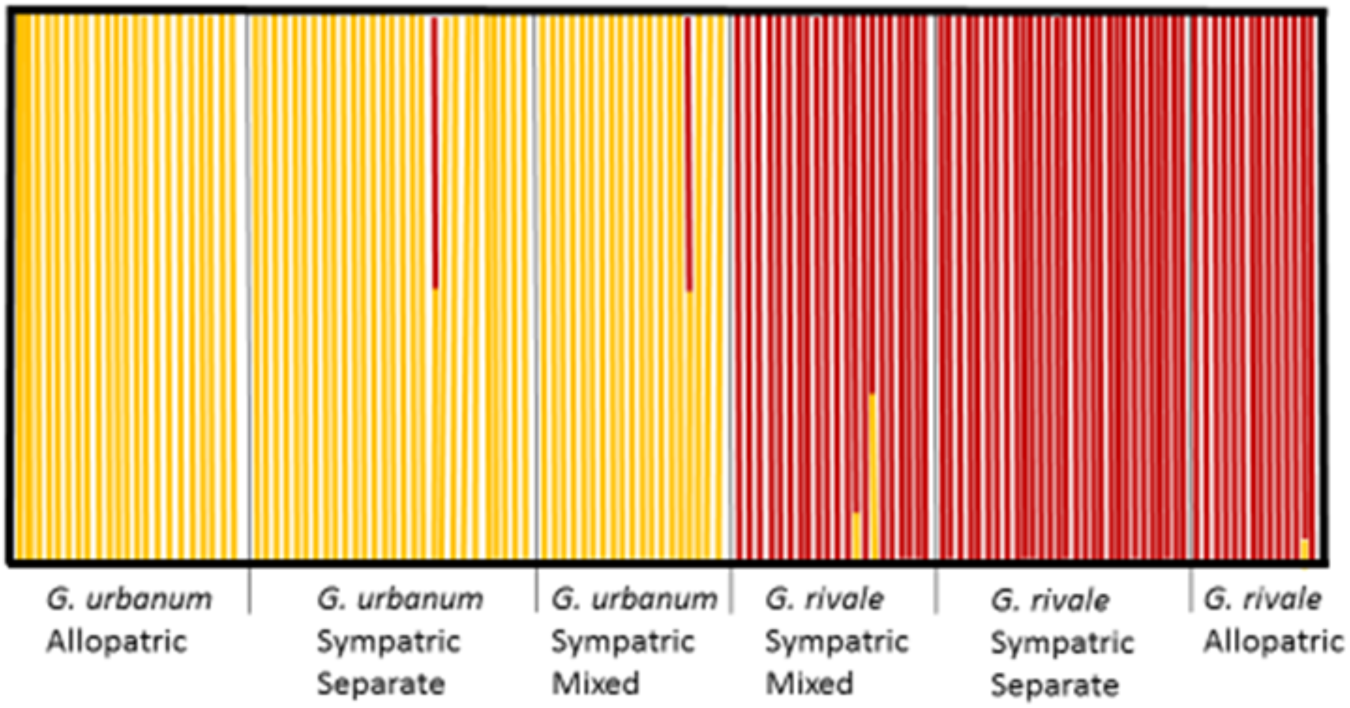
Results of fastSTRUCTURE (Raj et al. 2014) cluster analysis for K=2 based on 188 SNP loci present in all 133 *Geum* individuals scored in the allopatric and sympatric populations. Each bar represents one individual and shows the proportion of the genome from *G. rivale* (red) and *G. urbanum* (yellow).

**Figure 3.**
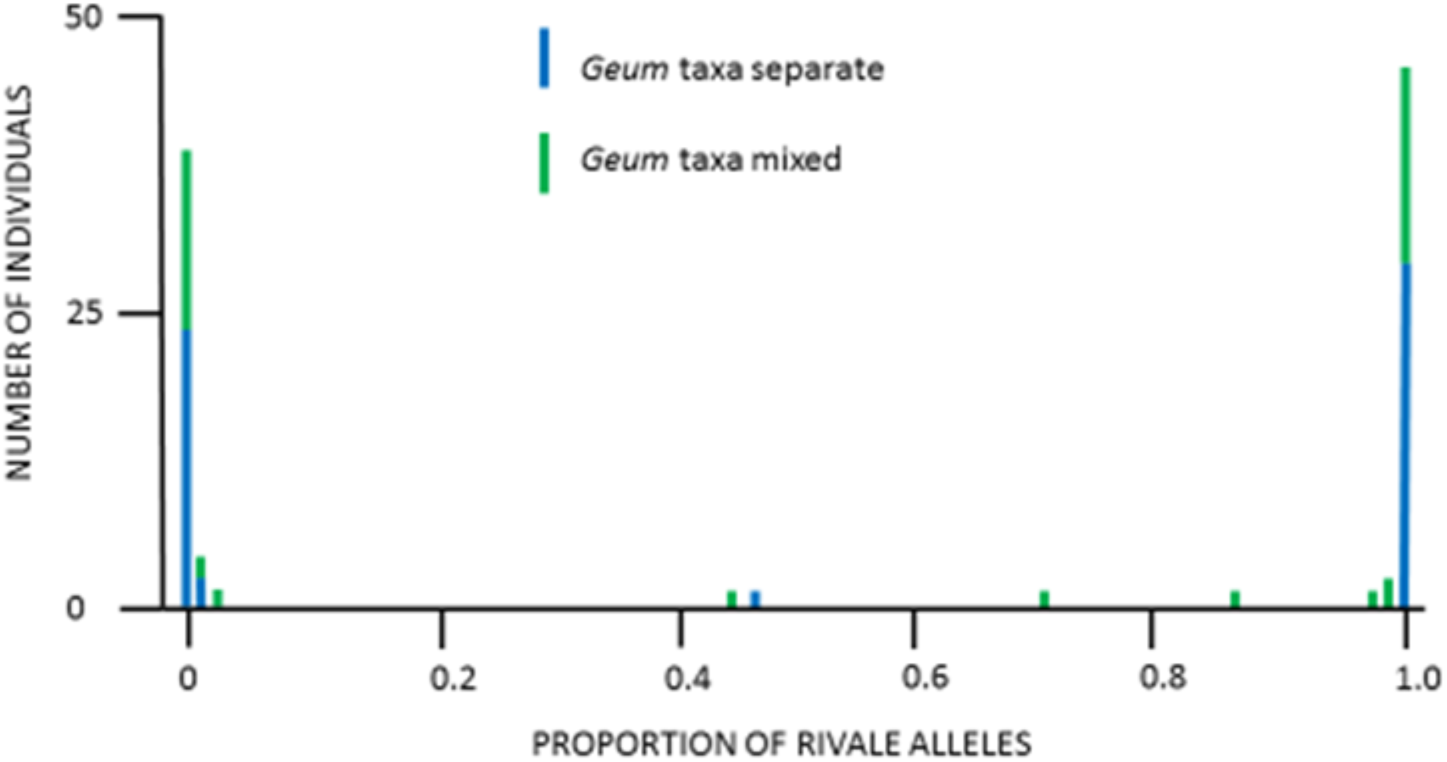
Distribution of hybrid index (proportion of *G. rivale* species-specific alleles) for 96 individuals within 39 sympatric populations where *G. rivale* and *G. urbanum* were either found alone (blue bars, ‘pure population’) or together (green bars, ‘mixed population’).

#### ii. Identifying hybrids and introgressed individuals using species-specific SNPs

To initially assess the ability of the species-specific SNPs to discriminate the species, we genotyped the 10 British allopatric *G. urbanum* samples that had not been used to generate species-diagnostic SNPs. 215 out of 220 diagnostic SNPs were homozygous for the *G. urbanum* allele in all individuals. Five SNPs were homozygous for the putative *G. rivale* allele in one or more of the 10 *G. urbanum* individuals, indicating that these SNPs were not fixed between species. They were removed from subsequent analysis of 96 individuals of *Geum* from the 39 sites sampled within the sympatric Berwickshire population. The mean number of SNPs successfully scored per individual was 207.3 (range 161 – 215 Figure S5). The variation in hybrid index (HI) in the Berwickshire population is illustrated in Figure 3.

The vast majority of individuals in the total sympatric sample (92 out of 96) have genotypes in which more than 97% of the species-specific SNPs are derived from only one of the two *Geum* species. Individuals possessing >97% *G. rivale* variants are hereafter regarded as *G. rivale*, while plants with >97% *G. urbanum* variants are classified as *G. urbanum.* The four remaining individuals with substantial proportions of variants from both species correspond to the four early generation hybrids identified in the fastSTRUCTURE clustering analysis. The correlation between the Q values calculated by fastStructure and the hybrid index calculated here is high (r=0.99) (see Figure S6 for comparison using 188 SNPs), in part because some SNPs (i.e., 61 of the 492 SNPs available for fastSTRUCTURE analysis) are common to both analyses. Early generation hybrids were significantly associated with ‘mixed’ rather than ‘pure’ sites within Berwickshire (Fisher’s exact test, P=0.044).

Neither of the two individuals possessing a hybrid index close to 0.5 are heterozygous at all SNPs, the situation anticipated if they were simple F1 hybrids (Table 3). Instead they are homozygous at a substantial fraction of the species-specific SNPs (15% and 18%), suggesting that they may have been derived by selfing of F1 hybrids. Two further individuals contain alleles derived predominantly from *G. rivale,* but additionally possess a substantial complement of *G. urbanum* alleles (29% and 12% respectively) (Table 5). On the basis of their complement of alleles alone, these plants are most likely to represent first and second generation backcrosses to *G. rivale* respectively. However, their origin must again be more complex, possibly involving selfing, because a substantial fraction of the *G. urbanum* alleles they possess (10% and 33% respectively) are present in homozygous form, whereas if they had been simple backcrosses, all *G. urbanum* alleles would have been present as heterozygotes.

**Table 5.** Genotypic composition at species-specific ddRAD loci for individuals classified as early generation hybrids (H) and backcrosses to *G. rivale* (BCR).

| Sample and classification | % '*rivale*' in genome | Homozygous '*rivale*' allele | Heterozygous | Homozygous '*urbanum*' allele | Total Loci scored |
| --- | --- | --- | --- | --- | --- |
| 29A (H) | 47.2 | 9 | 166 | 20 | 195 |
| 32C (H) | 45.3 | 7 | 132 | 22 | 161 |
| 26B (BCR) | 87.9 | 138 | 28 | 7 | 173 |
| 27B (BCR) | 70.8 | 95 | 113 | 6 | 214 |

Among the 92 plants from Berwickshire possessing a preponderance (>97%) of species-specific alleles from one taxon, 33 of the 47 individuals assigned to *G. rivale* and 26 of the 45 plants assigned to *G. urbanum* possessed from 1 to 10 alleles classified as specific to the alternate species (Fig. S7). These could represent alleles that have introgressed, or alternatively alleles present at low frequency in the focal species that have not been detected in the limited sample of allopatric genotypes used to identify species-specific SNPs. The maximum frequencies of putatively introgressed alleles are low (9.6% and 14.4% in *G. rivale* and *G. urbanum* respectively). Furthermore 76% are polymorphic in *G. rivale* only, 20% are polymorphic in *G. urbanum* only, and 4% are polymorphic in both species. These proportions are not significantly different from those found in the total sample of SNPs scored (P>0.05, Table 1). Therefore, misclassification of species-specific SNPs rather than introgression is the most likely explanation for the apparent presence of up to 3% admixture in the *Geum* genomes from sympatric populations.

With one exception, the putatively introgressed alleles discussed above were randomly associated with each other within each of the sympatric *G. rivale* and *G. urbanum* populations. However, two SNPs at radtags 20454 and 24791 respectively, and originally classified as specific to *G. rivale,* showed complete association in genotypic state within sympatric *G. urbanum.* The alleles were present at a mean frequency of 14.4% and distributed across six sites. In six individuals both alleles were present in homozygous form, while a seventh individual was heterozygous at both radtags. Since the SNPs involved are located on different scaffolds, these results may indicate the presence of a section of genome spanning the two scaffolds, possibly derived from *G. rivale,* which is segregating in the *G. urbanum* population.

### Analysis of Morphological Variation in Sympatric Populations

#### i. Correspondence between Phenotypic and Genotypic classification

Morphological measurements of eight floral and vegetative characters commonly used to distinguish *G. rivale* and *G. urbanum* were obtained from 87 of the 96 individuals from Berwickshire that had been scored for species-specific SNPs. The results of Principal Component Analysis based on these morphological data are illustrated in Figure 4. The first two principal components, accounting for 82.4% and 6.7% of the variation respectively, distinguish two major groups, with two individuals falling between these on the first PCA axis. Individuals classified as *G. rivale* (N=43) and *G. urbanum* (N=40) on the basis of species-specific SNPs fall clearly into the two major groups with +ve and –ve PCA1 values, respectively. The two individuals with intermediate PCA1 scores correspond to the early generation hybrids identified with clustering and species-specific SNPs. The two genotypes identified as complex backcrosses to *G. rivale,* on the basis of species-specific SNPs, group morphologically with individuals classified as *G. rivale* on the PCA plot.

**Figure 4.**
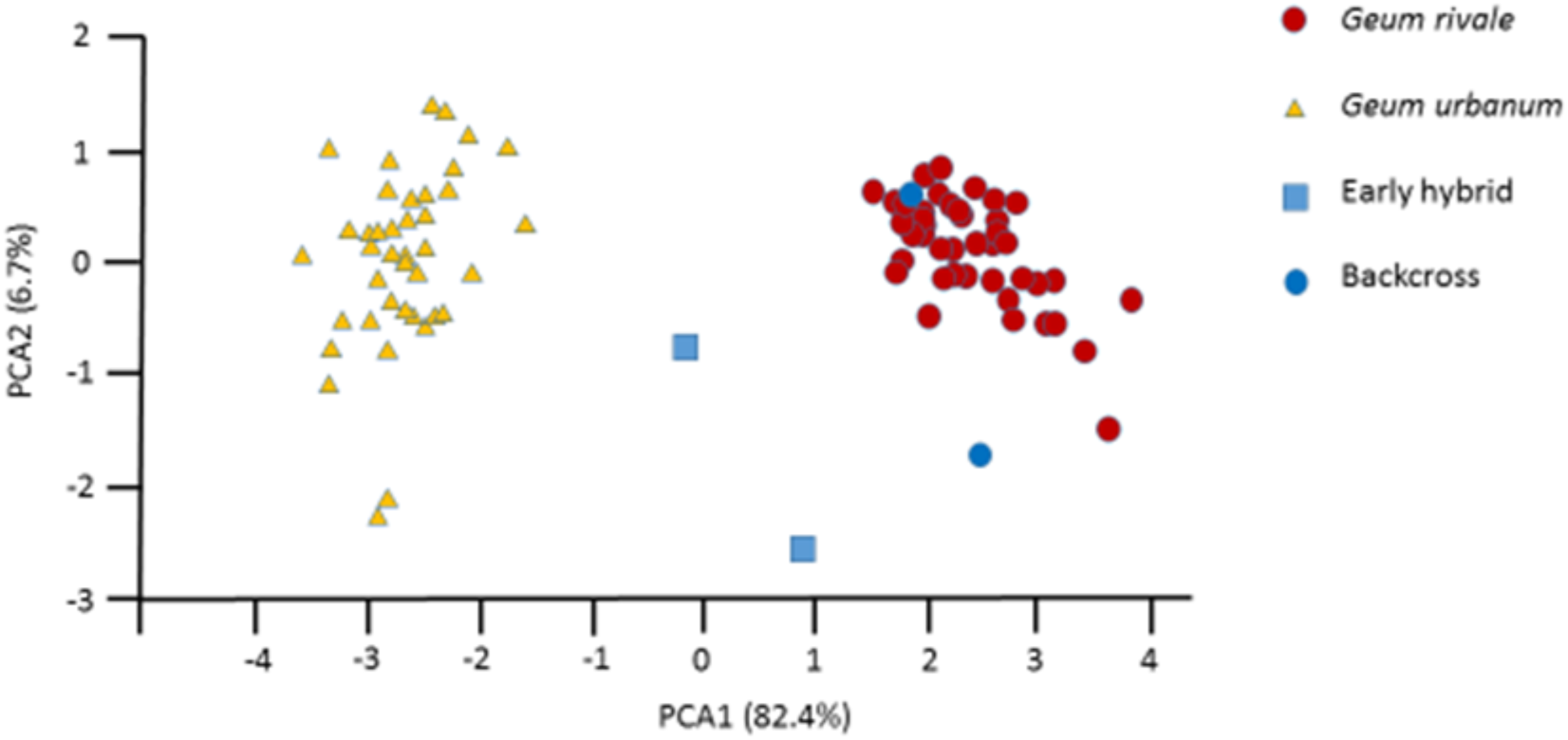
PCA analysis based on eight morphological characters measured in a common environment for a sample of 87 individuals of *Geum* sampled from sympatric populations. Genetic class (from Hybrid Index) of each individual (*G. rivale, G. urbanum,* early hybrid, or backcross to *G. rivale)* derived from an analysis of 215 species-specific markers is shown.

#### ii. Morphological Variation and Putative Introgression

To determine whether presence of putatively introgressed genetic material marked jointly by alleles at radtags 20454 and 24792 affected the morphology of *G. urbanum* (see above), data on the eight floral and vegetative characters measured in the sympatric *G. urbanum* population were used to generate the first two principal component scores describing morphological variation. Analysis of variance showed no significant effect of joint presence/absence of the alleles on PCA1, but a significant effect on PCA2 score (F_1,38_ = 8.67, P=0.005; Fig. 5). Individuals of *G. urbanum* jointly possessing these alleles showed a significantly larger flower angle than those lacking them (91.4 vs. 60.8 degrees, p=0.018). Greater flower angle is a feature characteristic of *G. rivale* and is most heavily weighted in PCA2.

**Figure 5.**
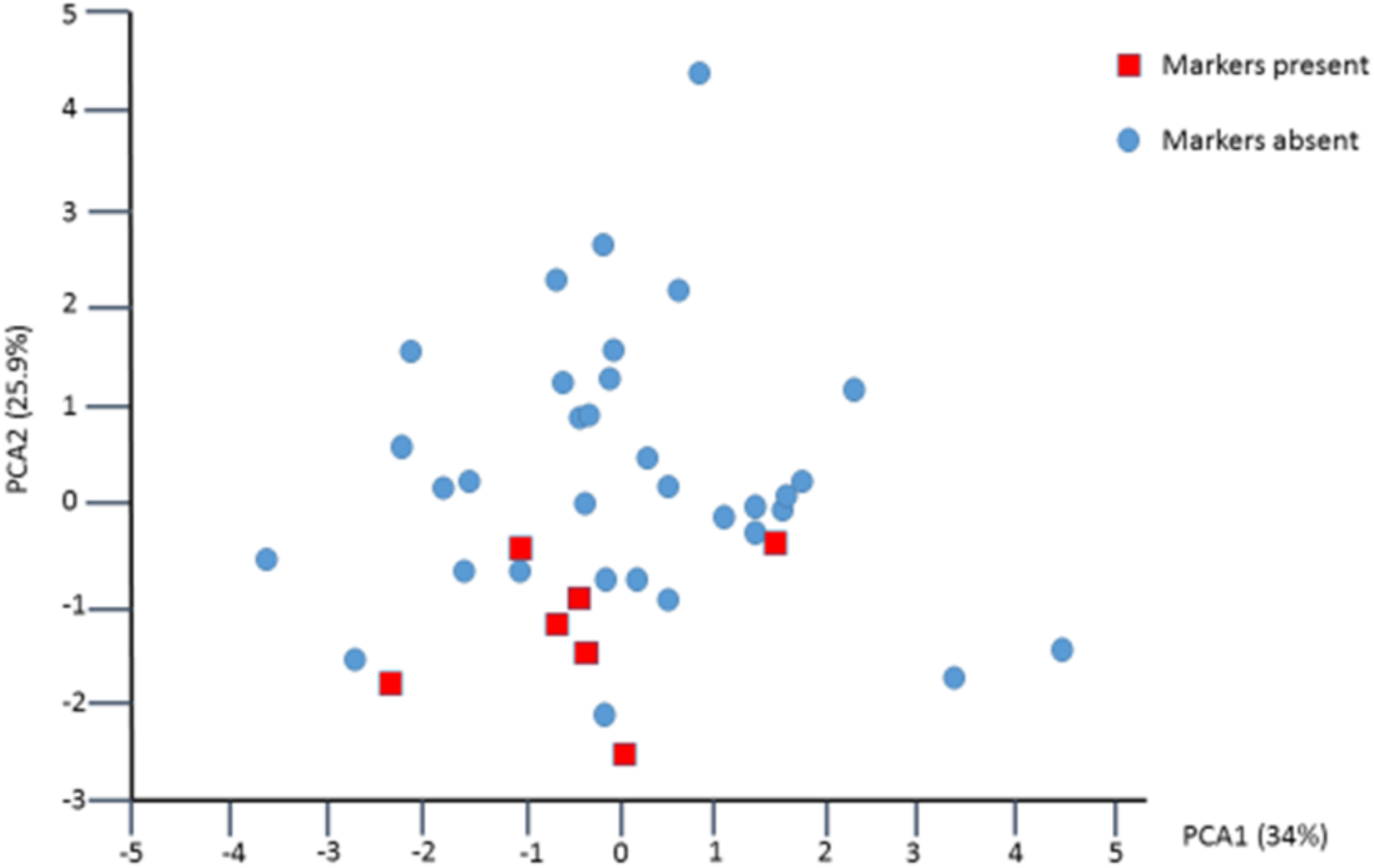
PCA analysis of 40 individuals of *G. urbanum* sampled from sympatric populations based on variation for eight morphological characters measured in a common environment. Red, square individuals jointly possess species-specific SNPs from *G. rivale* at radtags 20454 and 24791.

## DISCUSSION

We have developed a set of sequence based markers in two closely related non-model plant species with contrasting mating systems that share a polyploid history. Analysis of the markers across allopatric, and pure and mixed sympatric samples of *G. rivale* and *G. urbanum* indicate that the species are 2-3MY old, and that levels of introgression between the species, both before and after the period of recent sympatry, have been very small. While hybrid genotypes continue to be produced in contemporary populations, the proportion of introgressed loci across the genomes of the vast majority of individuals is much lower than 3%.

The development of sequence based markers, using next generation sequencing, poses technical problems in our *Geum* species. Their genomes are large (1.2 and 1.6 Gb in *G. urbanum* and *G. rivale* respectively) and although inheritance is disomic the species are ancient hexaploids. This means that there will generally be three closely related copies of every sequence, creating significant difficulties in distinguishing homologs from paralogs (Smedmark *et al.* 2003). We adopted the ddRAD technique that allows modulation of the number of RAD tags generated per genome to ensure adequate depth of coverage (Peterson *et al.* 2012). To maximize the chance of distinguishing homologs and paralogs we used paired end sequencing yielding long reads, and generated a draft genome of the largely homozygous inbreeding species against which to map our ddRADtag sequences. During the data analysis phase we employed very rigorous filtering of scaffolds and SNPs to exclude paralogs, which removed about three-quarters of scaffolds from our dataset. This filtering contributed to the development of a modest number of SNPs compared to comparable studies in other genera (e.g. *Populus*, Christe *et al.* 2016; *Betula*, Zohren *et al.* 2016), but that was nevertheless orders of magnitude higher than for previous microsatellite based studies; moreover, we developed sufficient SNPs to yield high precision estimates of introgression (e.g., 200 SNPs can estimate introgression of 0.5%). Furthermore our RAD data gave estimates of both inbreeding coefficients and population genetic differentiation that are congruent with results from microsatellite markers, providing confidence in the conclusions drawn from their analysis (Ruhsam *et al.* 2010, 2013).

Coalescent analyses of RAD data from allopatric populations, which contrast the fit of simple models of the species’ histories to polymorphism data, gave species’ age estimates of between 2 and 3 million years. This is considerably older than some estimates of time of divergence for other sister taxa that possess contrasting outcrossing and selfing mating systems. These range from 50 - 100kya for *Capsella grandiflora* and *C. rubella* (Brandvain *et al.* 2013) through 500-1100kya for *Mimulus guttatus* and *M. nasutus* (Aeschbacher *et al.* 2016) to 1Mya for *Arabidopsis lyrata* and *A. thaliana* (Tang *et al.* 2007). Our estimated age of the *Geum* species is consistent with the numerous morphological and physiological differences that are exhibited between them (Taylor 1997a, b). However despite their age, *G. rivale* and *G. urbanum* apparently exhibit few intrinsic hybrid incompatibilities (Ruhsam et al. 2013); these observations parallel those from *Drosophila* species, which show few post-zygotic incompatibilities for species of a similar age (Coyne & Orr 1989, 1997).

Coalescent analyses also show that low levels of effective introgression are likely to have occurred between the *Geum* species up to the time of isolation of allopatric populations, but there is no strong signal of introgression in a particular direction. For clarity, M measures the long-term effective migration, and accounts for reduced gene migration due to selection against hybrids (see below). Our estimate of the introgression rate between the *Geum* species (M=0.04 diploid genomes per generation) is an order of magnitude lower than that estimated for ancient introgression from inbreeding *M. nasutus* into outcrossing *M. guttatus* (M=0.1-1; Aeschbacher *et al.* 2016) and between outcrossing and selfing subspecies of *Clarkia* in their secondary contact zone (M=0.897-0.169; Pettergill & Moeller 2012). It is many orders of magnitude below contemporary estimates of unidirectional gene migration from the inbreeding rice *Oryza nivaria* to its outcrossing relative *O. rufipogon* (M=2.3 × 10^5^; Zhou *et al.* 2008).

Our coalescent analyses necessarily rely on simplifying assumptions. First, the blockwise site-frequency spectrum approach we adopted assumes a constant N_e_ and μ across blocks and so ignores effects of heterogeneity in mutation rate and the effects of background selection on linked sites, which could lead to a spurious signal of introgression (Ewing & Jensen 2016). Second, our models are drastic simplifications of the history of these species and, in particular, assume constant N_e_ in *G. rivale* and the ancestral population, which is unrealistic given the climatic fluctuations and the associated range shifts of European taxa during the Pleistocene. Following species divergence, either a change in N_e_ or introgression can alter the joint site-frequency spectrum (Chen 2012); by assuming N_anc_ = N_riv_, we potentially conflate changes in N_e_ with introgression, which, again, could lead to a spurious signal of introgression. Given these issues and the low estimates of introgression, we do not interpret the support of IM models over the div_2_ model as strong evidence for ancient long-term gene flow between the *Geum* species.

As a first test of more recent introgression in *Geum* we conducted comparisons of genetic diversity and differentiation between species in allopatry vs. sympatry. Introgression, which is only possible in sympatric populations, can potentially increase genetic diversity of the recipient species, and erode genetic differences between species. Hence, our comparisons of π and dxy between sympatric and allopatric populations can provide evidence for or against introgression. Relative genetic diversity was greater for *G. rivale* than *G. urbanum,* as expected based on their different mating systems (Charlesworth 2003). In addition, genetic diversity was approximately 9% higher for *G. rivale* in sympatry than in allopatry, but was equal between population types for *G. urbanum*. While this is consistent with introgression into *G. rivale*, but not *G. urbanum*, other factors could account for this result, including a greater effective population size of *G. rivale* in sympatry than in allopatry. Furthermore, genetic differentiation between species (dxy) was similar in sympatry and allopatry, providing no evidence for introgression.

As a second test of recent introgression we analysed the genomic composition of individuals from a region of sympatry in Berwickshire and compared them with samples from isolated, reference populations. The cluster analysis performed by fastSTRUCTURE (which did not rely exclusively on species-diagnostic SNPs), and the analysis based on a limited subset of species-diagnostic SNPs identified the same four individuals as early hybrids and backcrosses, and yielded similar genomic composition estimates that were congruent with morphological classifications. Critically, they suggest that the upper limit for the proportion of introgressed genome present in Berwickshire plants is ~3%. However, the real value for introgression is likely to be lower because the assumption that all our species diagnostic SNPs are alternately fixed between the species is unlikely to be true.

Although our overall results indicate very little recent genetic exchange, one intriguing result provides tentative evidence for introgression from *G. rivale* to *G. urbanum*. Two SNP variants located on different scaffolds that were classified as specific to *G. rivale* in allopatry show co-segregation within the sympatric *G. urbanum* population. Moreover, *G. urbanum* individuals possessing these alleles show a significantly greater flower angle, a characteristic of *G. rivale*. A possible explanation is introgression of a chromosome segment from *G. rivale* into *G. urbanum* that is now polymorphic within the sympatric population. Given the absence of evidence for backcrossing of hybrids to *G. urbanum* (Ruhsam *et al.* 2011, 2013), such introgression could only arise through the establishment, following hybridisation, of a self-fertilising lineage that largely retains genetic material from *G. urbanum*. Previous work has shown that F1 hybrids are self-compatible and a proportion may be capable of autopollination, so this scenario, though unlikely, is not impossible (Ruhsam *et al.* 2013).

Despite the reproductive barrier that adoption of a selfing mating system naturally imposes (Martin & Willis 2007; Briscoe Runquist *et al.* 2014; Brys *et al.* 2014), the widespread occurrence of the F1 hybrid *G. intermedium* throughout the area of sympatry of *G. rivale* and *G. urbanum* in Berwickshire and elsewhere (Preston *et al.* 2002), and our own detection of early generation hybrids, indicates that low levels of introgression are not caused by lack of opportunity for hybridisation. Previous studies have shown that where hybridisation occurs, a wide array of fertile recombinant offspring are produced both by backcrossing with *G. rivale* and by self-fertilisation and that under benign conditions these show little evidence of intrinsic genetic incompatibility (Ruhsam *et al.* 2011, 2013). Thus *G. rivale* and *G. urbanum* join a growing list of sister species which readily produce F1 hybrids and viable recombinant offspring, yet show little or no signal of introgression (e.g. *Encelia* (Kyhos *et al.* 1981); *Quercus*, *Arctostaphylos*, *Iris* (Nason *et al.* 1992); *Rhododendron* (Milne *et al.* 2003); *Begonia* (Twyford *et al.* 2015); *Populus* (Christe *et al.* 2016); *Bombina* (Nürnberger *et al.* 2016).

In many of the above examples ecological selection against recombinants has been invoked to explain the maintenance of species distinctions. There are good grounds for proposing the same explanation to account for lack of introgression in *Geum*. Since their initial divergence the *Geum* species have evolved marked differences in pollination syndrome, phenology, vegetative morphology and many aspects of physiology including tolerance of shade and waterlogging (Waldren *et al.* 1988; Taylor 1997a, b). Therefore multiple ecologically important characters could potentially be the targets of selection against recombinant individuals in natural populations (Nosil *et al.* 2009). Selection may be particularly effective in the self-fertilising recombinants that are required to allow introgression from *G. rivale* into *G. urbanum*. Here reduced effective recombination rates will lead to correlated selective effects across the genome, and deleterious recessive mutations present as genetic load in genome segments derived from outcrossing *G. rivale* will be exposed (Hu 2015). Given our ability readily to artificially cross the *Geum* species, and produce large numbers of later generation hybrids, there is now the potential experimentally to test the hypothesis that strong ecological selection maintains species integrity using appropriate field experiments.

## Acknowledgements

This research was funded by BBSRC grant Bb/J006580/1 to RAE. We are very grateful to Barbara Giles, Cecile Bacles, Olivier Lepais, Hjörtur Porbjörnsson, and Jorge Paiva who provided non-UK samples for this project. Beate Nürnberger aand Alex Tywford provided comments that improved this paper.

## Author contributions

C.Y.J, K.G. and R.A.E. conceived the study; C.Y.J., F.T., M.T. and R.A.E. gathered data; C.Y.J., K.L., F.T. and R.A.E. analysed data; C.Y. J., K.L. and R.A.E. wrote the manuscript with input and revisions from all co-authors.

## Data accessibility

DNA sequences: will be made available on European Nucleotide Archive before publication

Assembled genome: will be made available (University of Edinburgh link) before publication

Morphological data: will be made available on Dryad before publication

Scripts: will be made available on github.com before publication

## Conflict of interest

None declared.

## SUPPLEMENTARY MATERIAL

### i. Development of Draft Genome for *G. urbanum*

Reads were generated from three Illumina libraries: a 151 paired end library with a mean insert size of 250 (sequenced on Hi seq 2500 (v1 chemistry) to give 144 million read pairs), a 151 paired end library with a mean insert size of 450 (sequenced on Hi seq to give 134 million read pairs), and a 251 paired end library with a mean insert size of 450 (sequenced on Illumina Mi seq (v2 chemistry) to give 11.9 million read pairs). Illumina adapters were removed and reads quality trimmed using cutadapt version 1.3 (Marcel, 2011) with the option “-q 30”. Trimmed reads were assembled into contigs and scaffolds using MaSuRCA version 2.0.3.1; MaSuRCA (Zimin et al., 2013) was run using the command line options:

GRAPH_KMER_SIZE=auto, USE_LINKING_MATES=1, LIMIT_JUMP_COVERAGE = 60, ovlMerSize=30, cgwErrorRate=0.25, ovlMemory=4GB, KMER_COUNT_TH RESHOLD = 1, NUM_THREADS= 64,JF_SIZE=23000000000, DO_HOMOPOLYMER_TRIM=0.

### ii. Sample Preparation, Library Preparation, and PCR Amplification for ddRAD

For each individual sample, 35ul of genomic DNA (250ung) was digested in a total volume of 50 ul, comprising 1.0 ul SphI-HF (20u), 1.0 ul EcoRI-HF (20u), 5.0 ul 10x Cutsmart buffer (New England Biolabs, UK) and 8.0 ul dH2O for a total of 3 hours, at 37°C. The reaction was purified by the addition of Ampure XP beads at a ratio 1.8 x beads: 1x DNA rather than by heat denaturation, to maintain the integrity of the EcoRI generated overhangs.

Samples were eluted in 25ul EB (Qiagen). Unique barcoded EcoRI P1 adapters (Peterson et al. 2012) and a generic SphI P2 adapter (Table S3) were ligated to the complementary overhangs of the DNA fragments generated by the EcoRI:SphI double digest reaction. Each ligation reaction was carried out in a total volume of 40ul, consisting of 25ul DNA, 2.0 ul P1 adapter (4uM), 2.0 ul P2 adapter (4uM), 4.0 ul 10x T4 Ligase buffer (New England Biolabs, UK) 1ul Quick Ligase (New England Biolabs, UK), and the reaction was incubated at 23°C for 30 min followed by heat denaturation at 65 °C for 15 min.

After ligation the reactions were pooled into libraries containing a maximum of 24 samples, each barcoded with a unique EcoRI PI adapter sequence. The pooled libraries were then subjected to two rounds of Ampure XP bead clean up, at a ratio of 1x Ampure XP beads: 1 x DNA and eluted in a volume of 30ul EB (Qiagen). Individual libraries were size selected by BluePippin Size selection system, 2% Precast Agarose gel cassette (sage science) using the 'Tight' collection mode set to a target size of 300bp. The success of size selection was assessed by Bioanalyzer DNA HS chip (Agilent Technologies).

For each ddRAD library, PCR amplification was carried out in 2 independent 50 μL reactions consisting of 20 ul size selected DNA, 25ul 2x Phusion Master Mix (New England Biolabs, UK), 2.5 ul DMSO, and 1ul ddRAD P1 PCR primer (10 nm), and a barcoded P2 primer (10 nm) that introduced a 8bp unique index during amplification (Table S3), using the following cycling parameters: 98 °C for 30 s followed by 12 cycles of 98 °C for 10 s and 72 °C for 60 s.

PCR products were then pooled, and purified by Ampure XP beads at a ratio of 0.8 x beads: 1x DNA, and the mean insert size of the individual libraries was assessed by Bioanalyzer DNA HS chip, (Agilent Technologies), and were quantified by qPCR on an Illumina Eco instrument using the Kapa Library Quantification Kit.mA final super pool was created consisting of equimolar individual library PCR reactions and the quantification of the pool was measured by qPCR.

## SUPPLEMENTARY TABLES

**Table S1.**
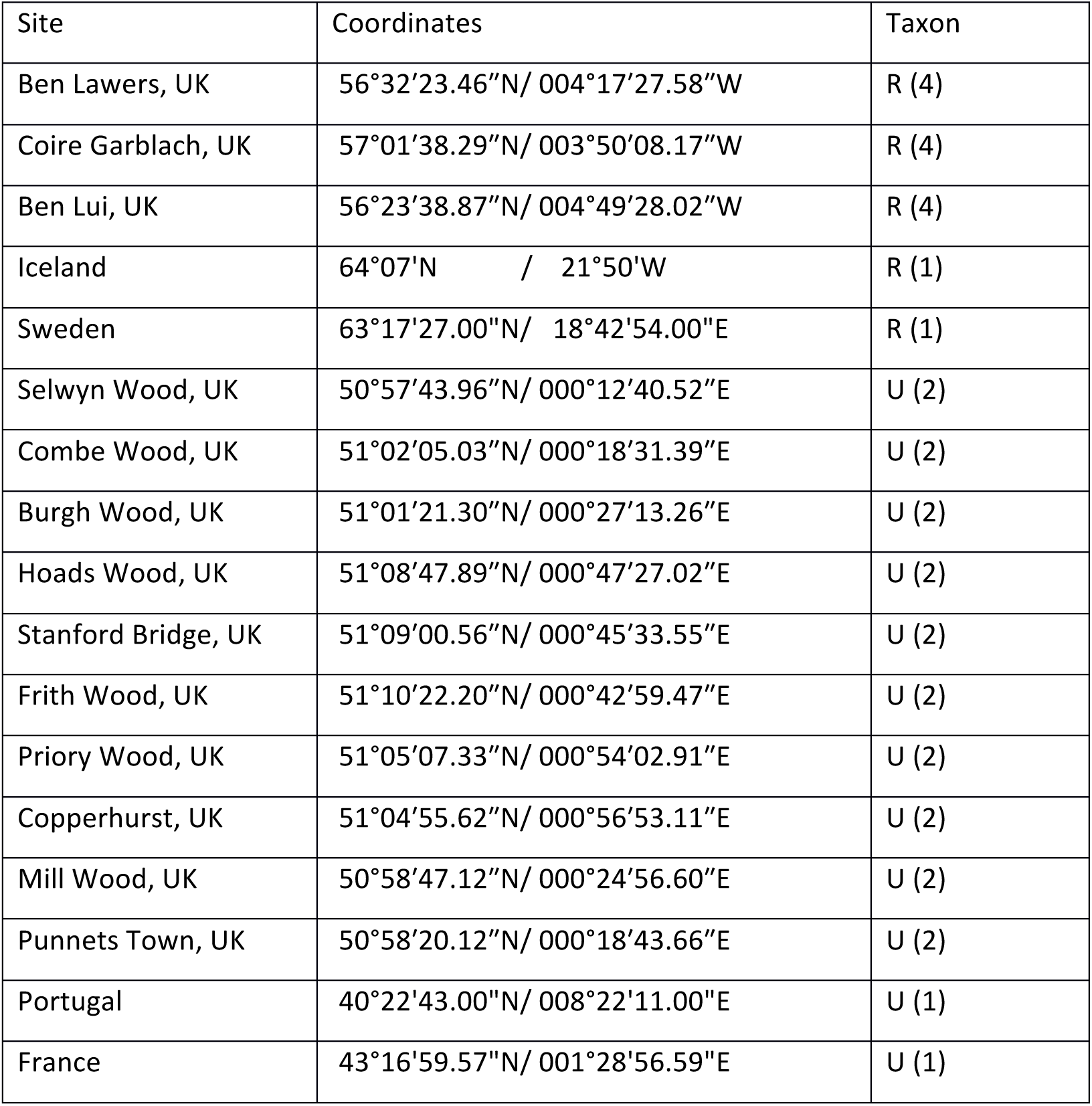
Table S1. Sampling Locations for allopatric populations of *Geum rivale* (R) and *G. urbanum* (U). Numbers in brackets = sample size.

**Table S2.**
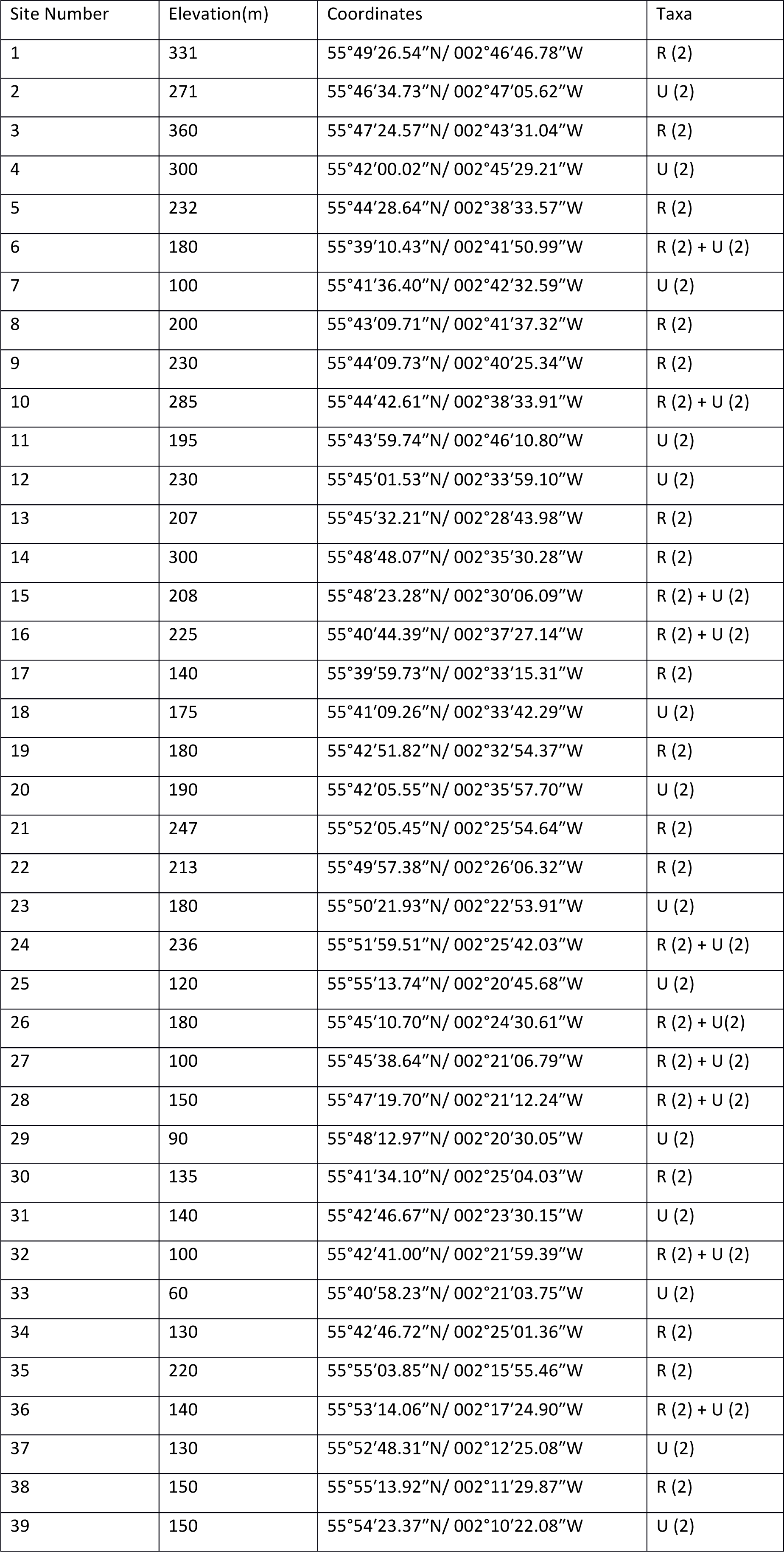
Sampling locations for pure and mixed populations of *Geum rivale* (R) and *G.rivale* (U) 924 within an area of geographic sympatry in Scotland. Numbers in brackets = samplesize.

**Table S3.**
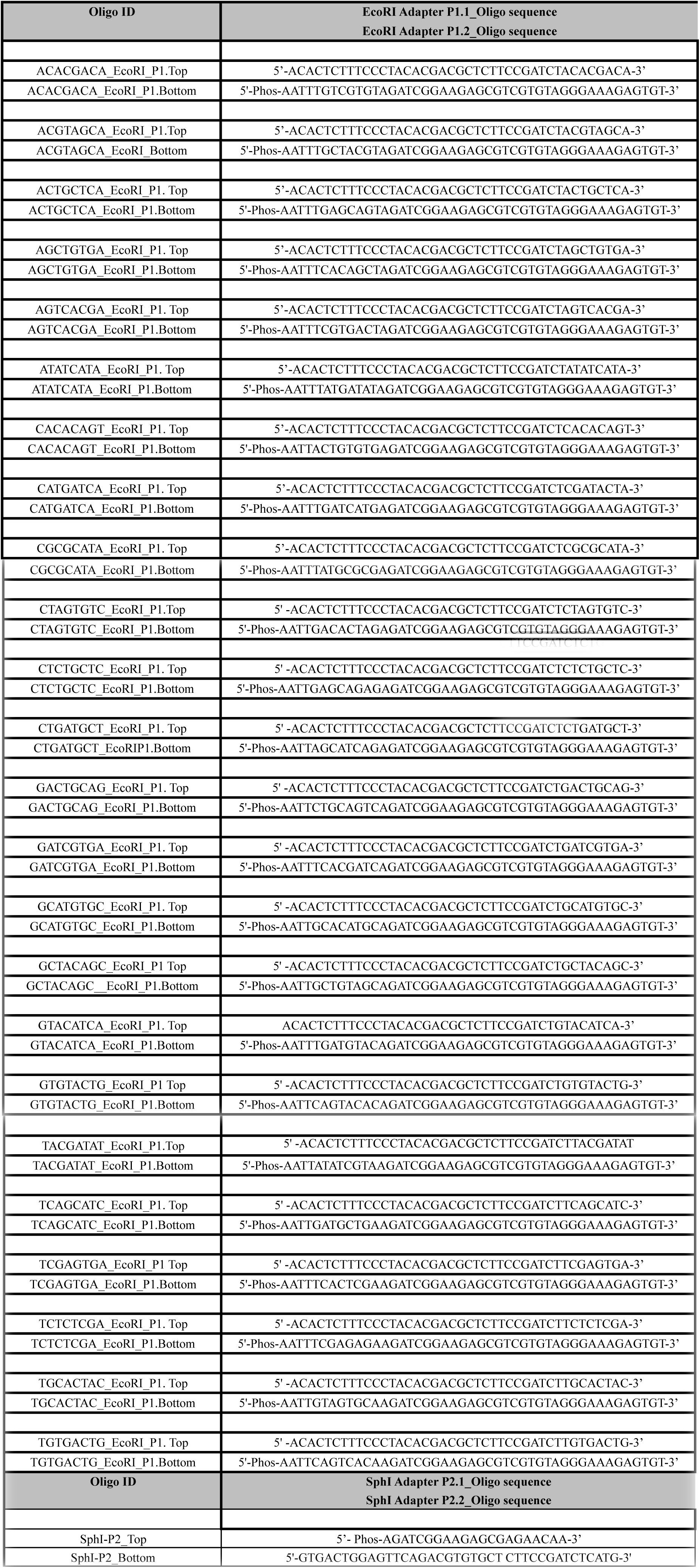
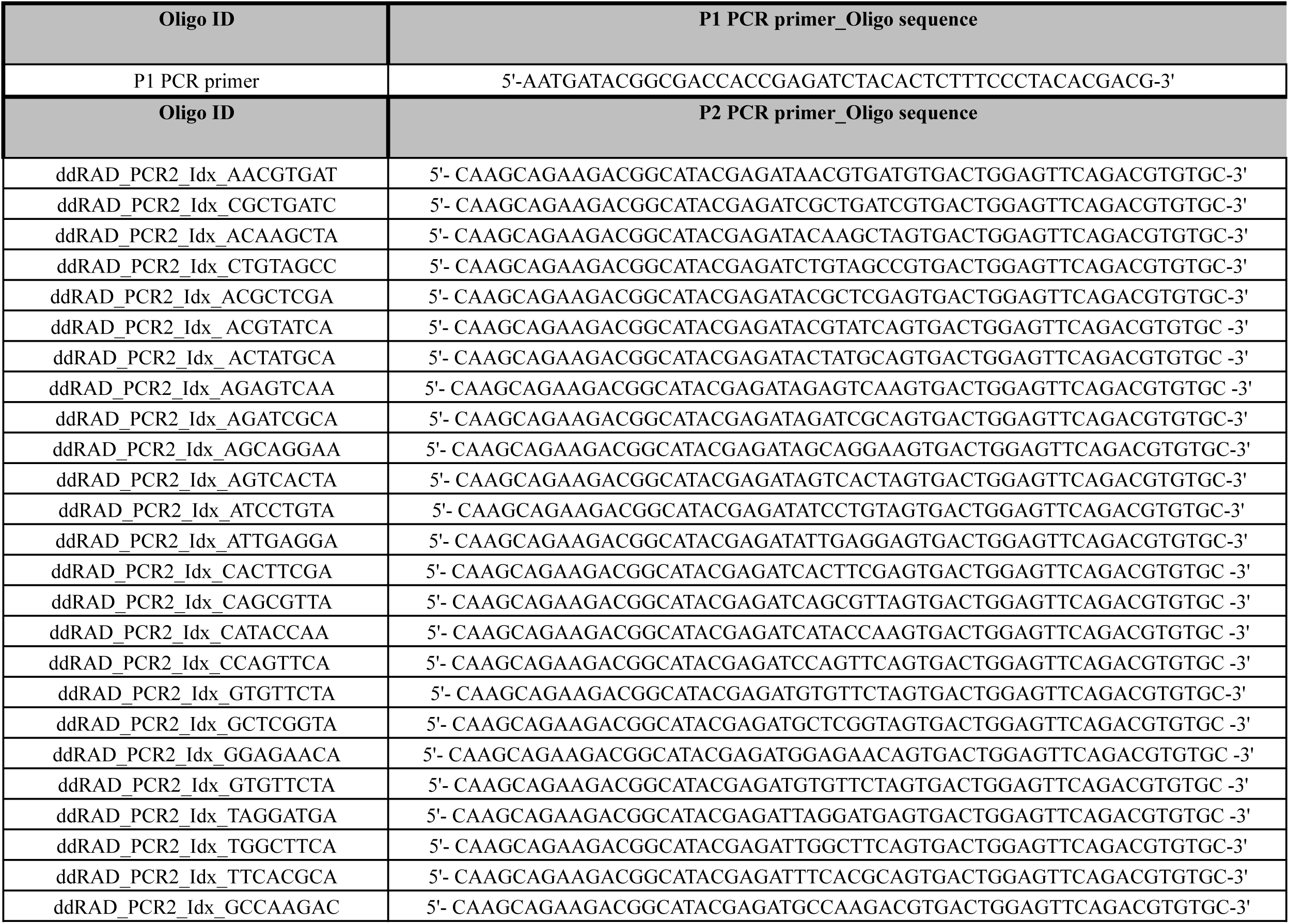
EcoRI P1 adapters and generic SphI P2 adapter; P1 and P2 primers used in ddRAD amplification

**Table S4.** F_IS_ estimates for ‘allopatric’ UK *G.rivale* and *G.urbanum* populations using data data not filtered for paralogs.

|  | Unfiltered for paralogs |  |
| --- | --- | --- |
| Population by species | Number of SNPs | $F_{IS}$ |
| *G. rivale* |  |  |
| Ben Lawers | 2531 | -0.0082 |
| Coire Garblach | 2605 | 0.0384 |
| Ben Lui | 2664 | 0.0423 |
| *G. urbanum* |  |  |
| Priory Wood | 551 | 0.6216 |
| Mill Wood | 531 | -0.0697 |
| Burgh Wood | 542 | 0.5913 |
| Hoades Wood | 451 | 0.5100 |
| Punnetts Town | 495 | 0.4636 |
| Frith Wood | 507 | 0.6134 |
| Stanford Bridge | 535 | 0.6112 |
| Combe Wood | 536 | 0.6110 |
| Copperhurst | 509 | 0.5874 |
| Selwyn Wood | 166 | -0.4759 |

**Table S5:** Polymorphism counts and numbers of DNA blocks for coalescent analysis for three individual *G.rivale* samples from UK allopatric populations.

| Sample | Heterozygosity | No. SNPs private to *G. rivale* | No. SNPs private to *G. urbanum* | No. SNPs shared by both species | No. alternately fixed SNPs | No. DNA blocks |
| --- | --- | --- | --- | --- | --- | --- |
| Ben Lui 1 |  | 250 | 88 | 0 | 1184 | 668 |
| Ben Lui 4 |  | 299 | 86 | 1 | 1170 | 672 |
| Ben Lawers 5 |  | 179 | 88 | 0 | 1189 | 649 |

**Table S6.** Parameter estimates under models analysed for sample pairs involving the samples: Ben Lui 1, Ben Lui and Ben Lawers 5.95% Confidence intervals provided in parentheses. See FigureS4 for plots of 95% CI’s.950

|  | Model |  |  |
| --- | --- | --- | --- |
| | $\text{div}_2$ | $\text{IM}_{r \rightarrow u}$ | $\text{IM}_{u \rightarrow r}$ |
| **Ben Lui 1** |  |  |  |
| $N_{\text{anc}} = N_{\text{rivb}}$ | $1.03 * 10^5$ | $1.01 * 10^5$ | $6.42 * 10^4$ |
| $N_{\text{urb}}$ | $3.46 * 10^4$ | $3.23 * 10^4$ | $3.29 * 10^4$ |
| $t$ (years) | $2.36 * 10^6$ | $2.41 * 10^6$ | $3.03 * 10^6$ |
| $M$ | | 0.02 (0.00 – 0.18) | 0.08 (0.06 – 0.11) |
| **Ben Lui 4** |  |  |  |
| $N_{\text{anc}} = N_{\text{rivb}}$ | $1.30 * 10^5$ | $1.24 * 10^5$ | $1.11 * 10^5$ |
| $N_{\text{urb}}$ | $3.77 * 10^4$ | $3.09 * 10^4$ | $3.38 * 10^4$ |
| $t$ (years) | $2.22 * 10^6$ ($2.01 * 10^6 - 2.43 * 10^6$) | $2.35 * 10^6$ | $2.48 * 10^6$ |
| $M$ (mean, 95% CI's) | | 0.04 (0.005 - 0.191) | 0.04 (0.007 - 0.101) |
| **Ben Lawers 5** |  |  |  |
| $N_{\text{anc}} = N_{\text{rivb}}$ | $7.41 * 10^4$ | $7.34 * 10^4$ | $5.11 * 10^4$ |
| $N_{\text{urb}}$ | $3.54 * 10^4$ | $3.40 * 10^4$ | $3.38 * 10^4$ |
| $t$ (years) | $2.60 * 10^6$ | $2.64 * 10^6$ | $3.10 * 10^6$ |
| $M$ (mean, 95% CI's) | | 0.01 (0.00 – 0.103) | 0.02 (0.002 - 0.052) |

## SUPPLEMENTARY FIGURES

**Figure S1:**
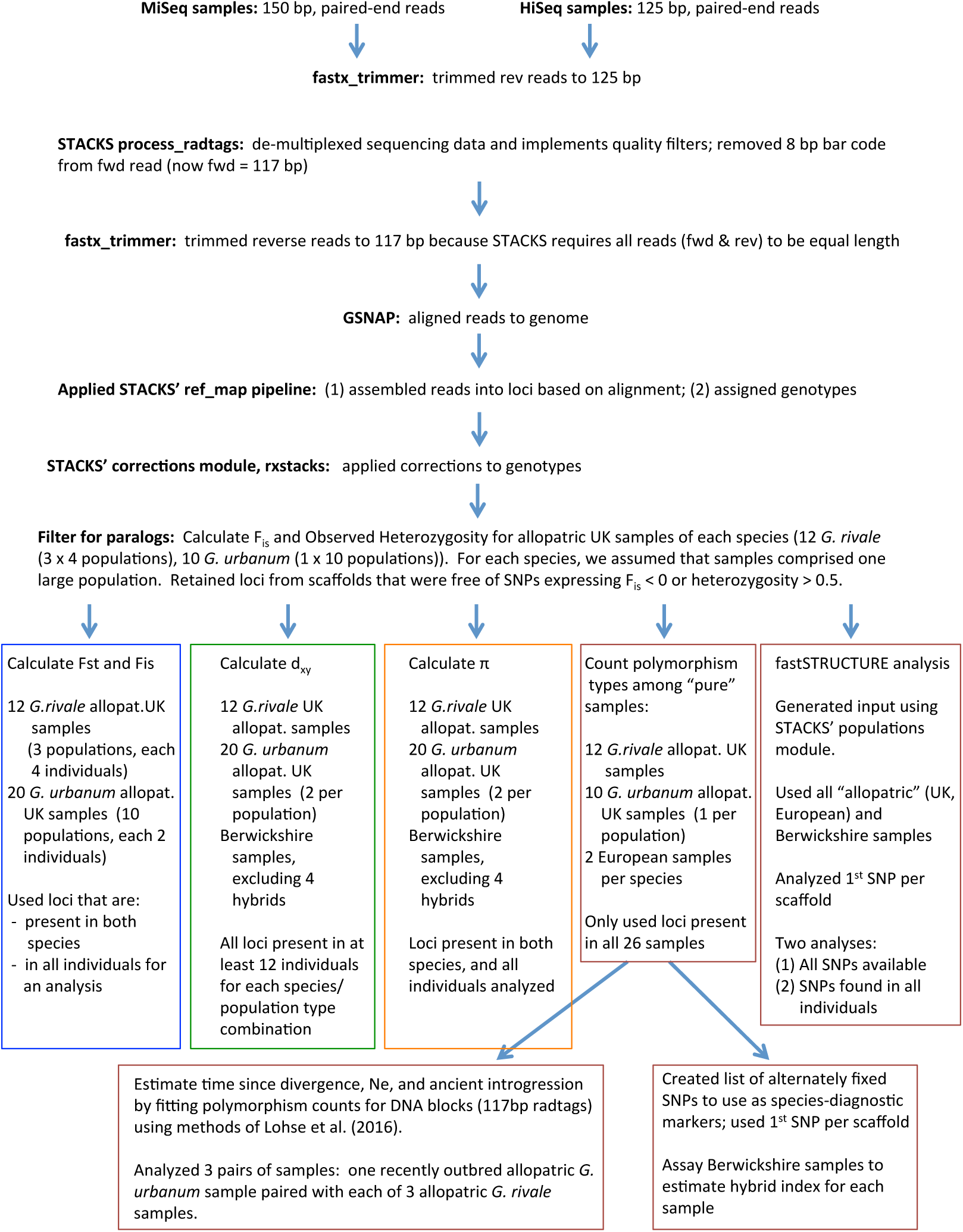
Bioinformatics pipeline for analysis of *Geum* ddRAD sequences.

**Figure S2.**
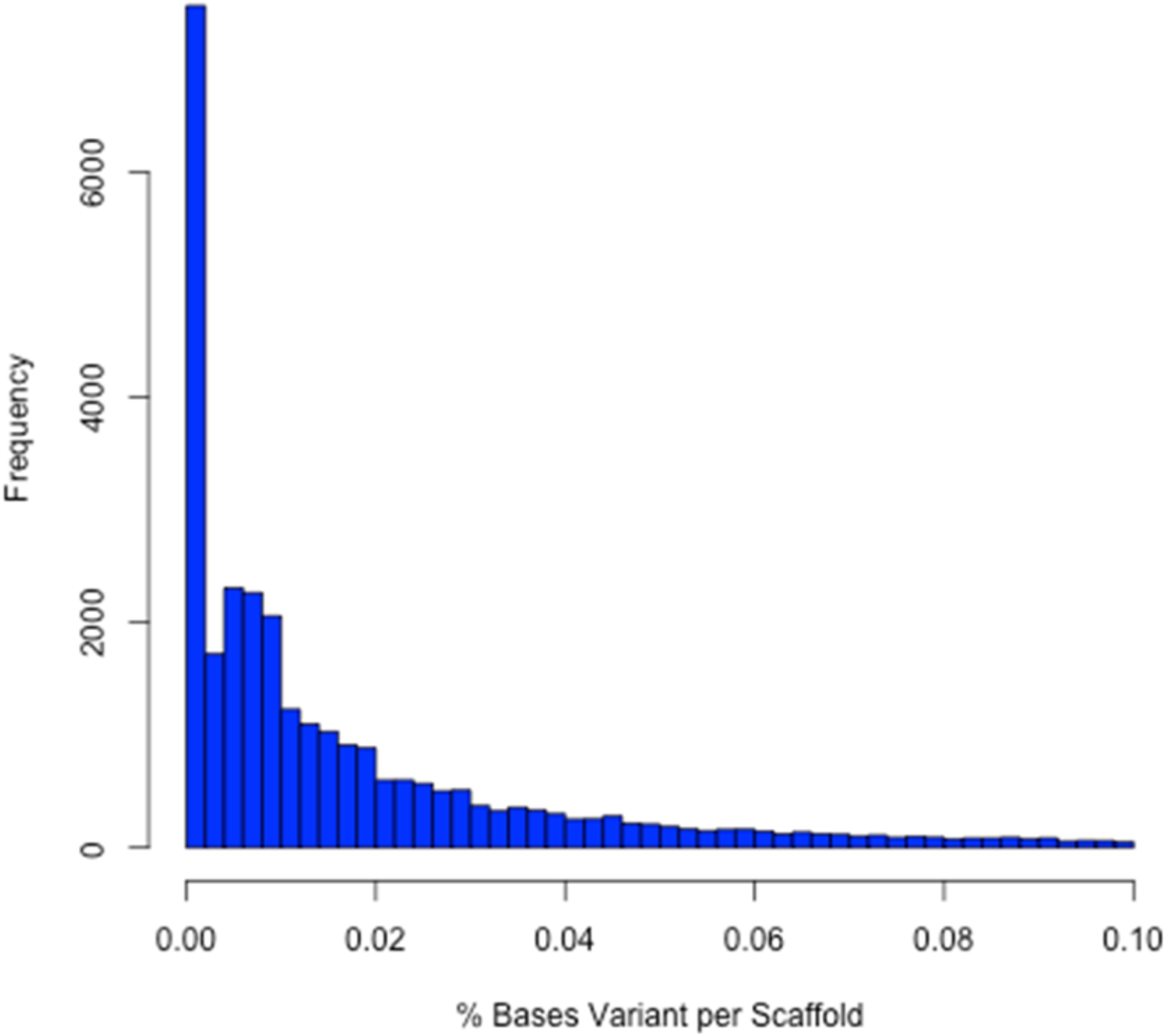

**Figure S3.**
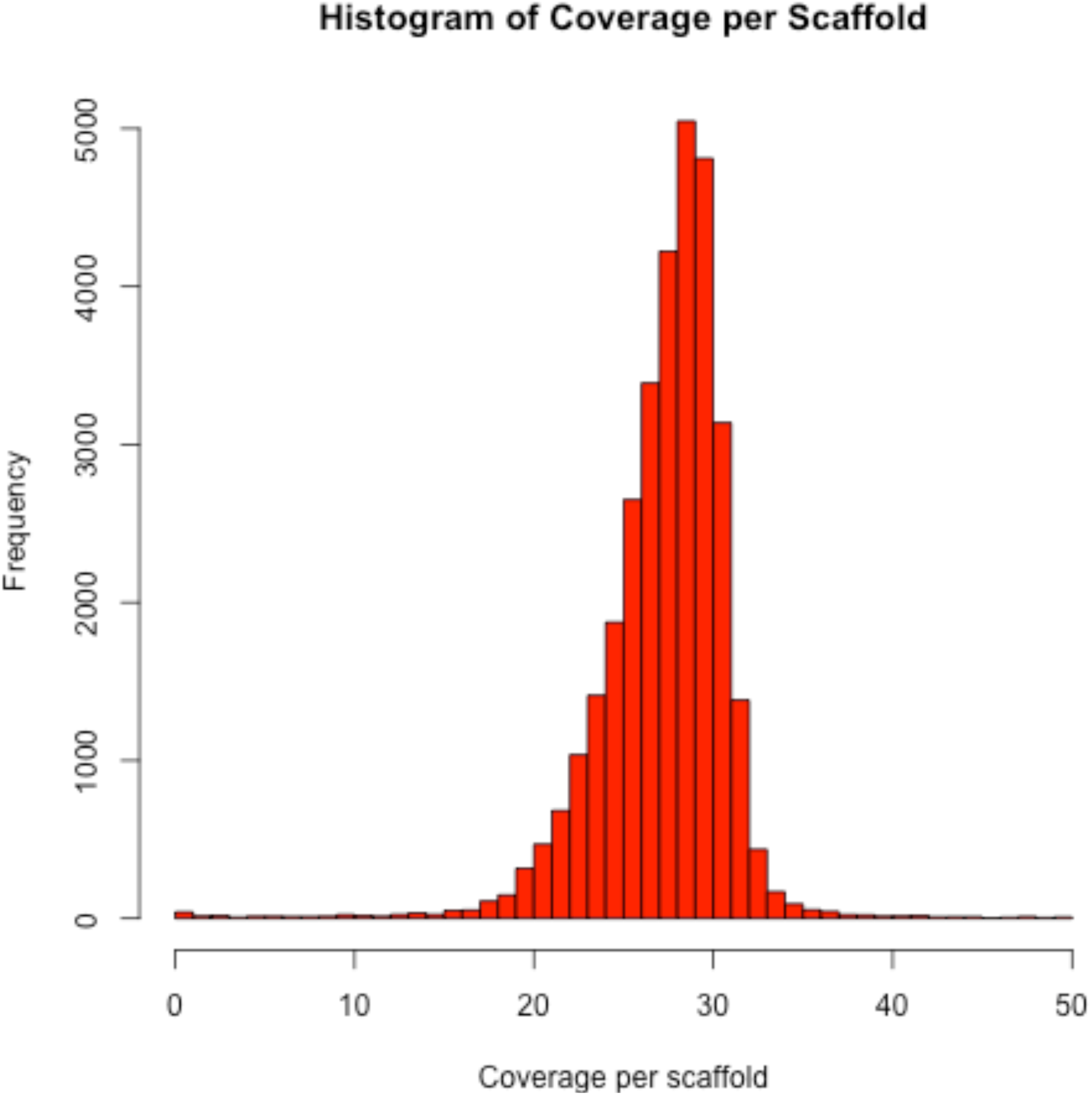

**Figure S4:**
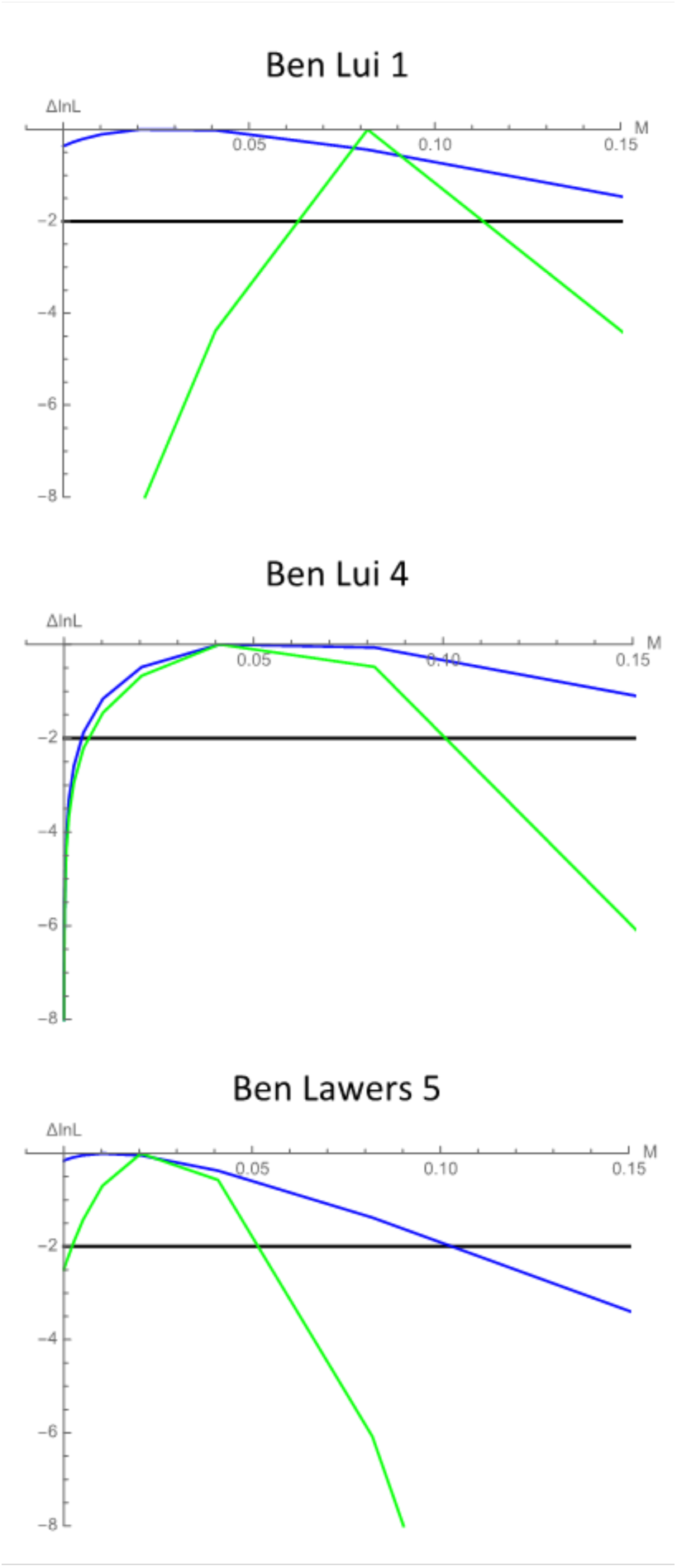
Log-Likelihood plots for estimates of M for three pairs of *Geum* samples, each involving a different *G. rivale* individual (Ben Lui 1, Ben Lui 4, Ben Lawers 5). Blue and green lines represent log-likelihoods of M for models IM_r→u_ and IM_u→r_, respectively. 95% CI’s are bounded where these lines intersect the dark horizontal line.

**Figure S5.**
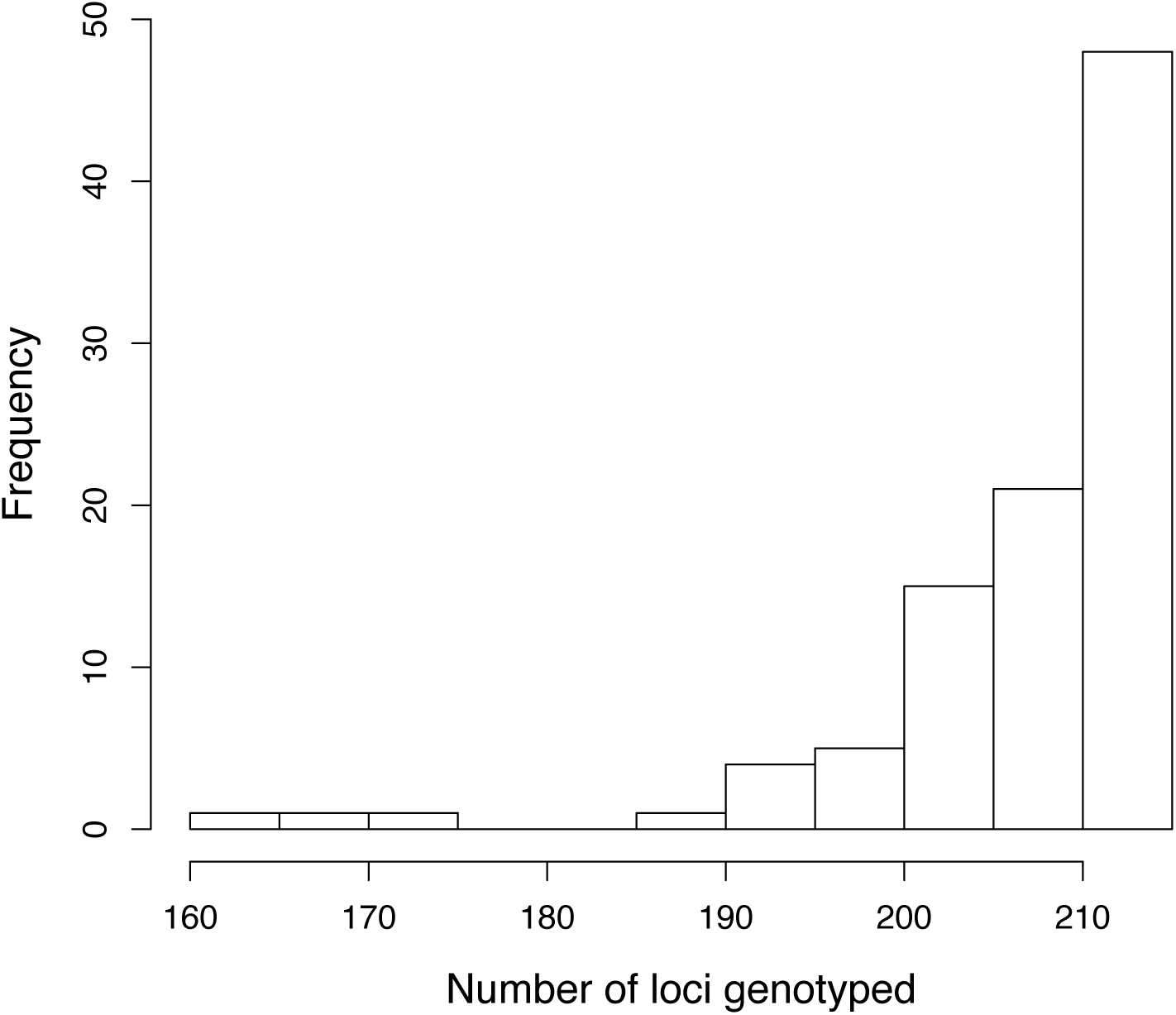
Number of species-specific SNPs scored per individual in a sample of 97 *Geum* plants from 39 populations in Berwickshire.

**Figure S6.**
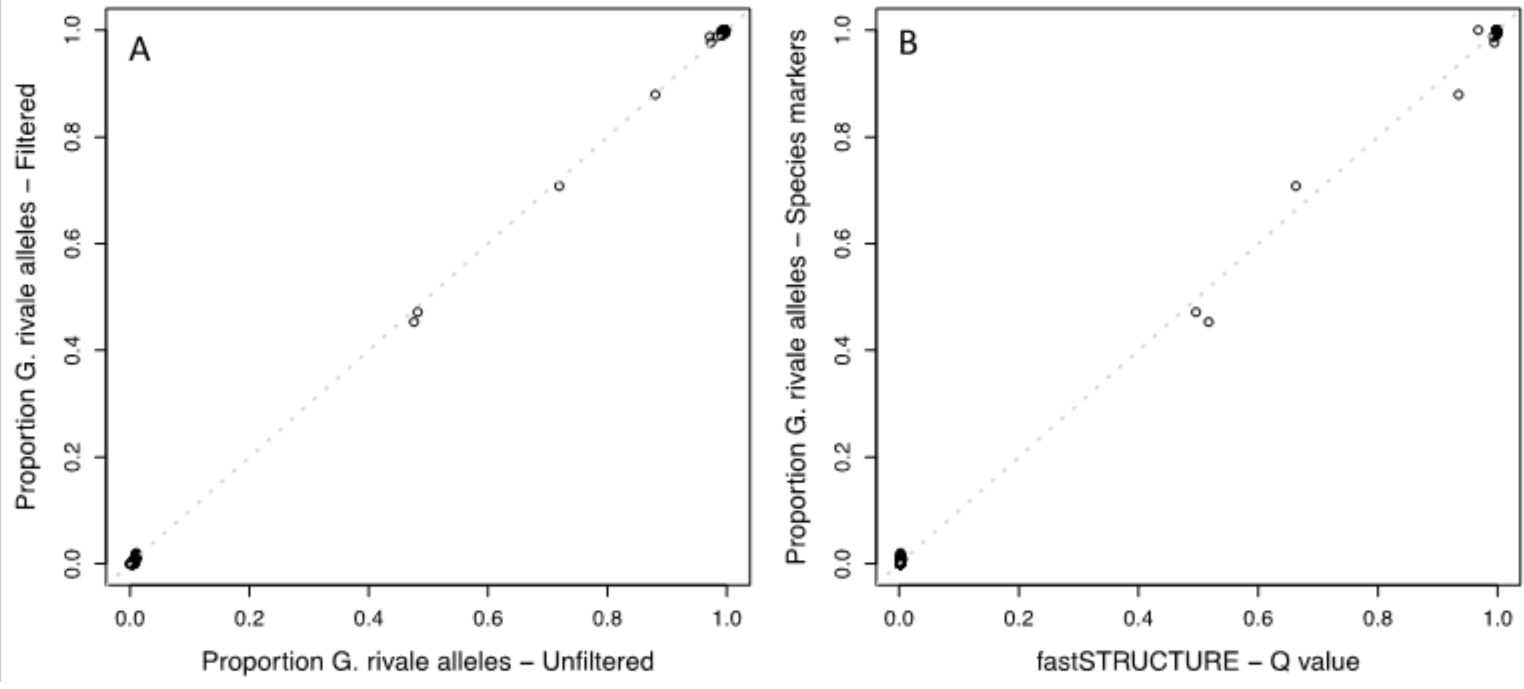
Genomic composition estimates for Berwickshire samples derived from paralog-filtered species-diagnostic SNPs (y-axis in both panels) compared to analogous measures in two alternate analyses: (A) species-diagnostic SNPs from data unfiltered for paralogs, and (B) Admixture proportion (Q value) calculated by fastSTRUCTURE analysis that required loci to be present in all individuals (188 SNPs).

**Figure S7.**
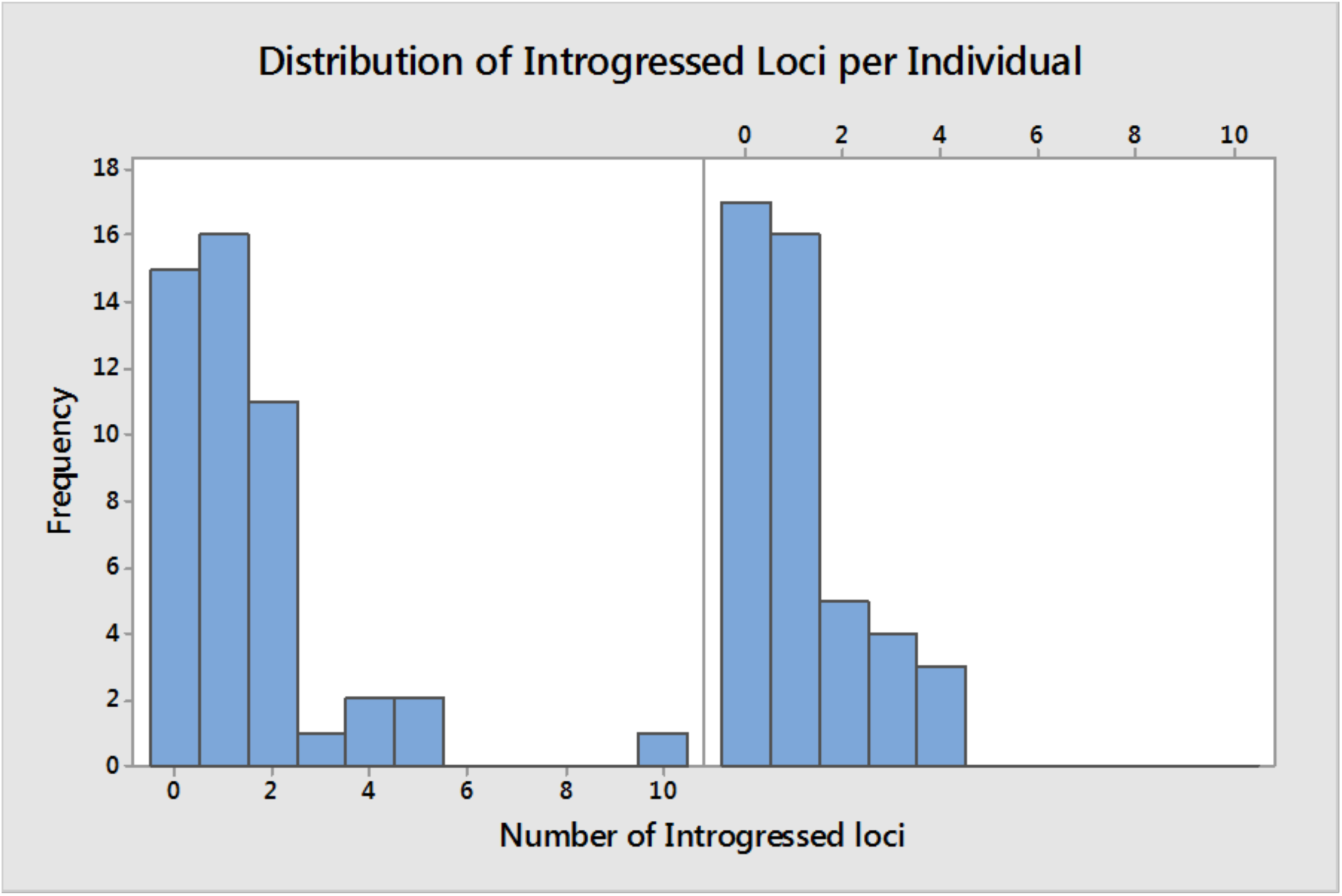
Number of putatively introgressed loci per individual in individuals classified as *G. rivale* (left panel) and *G. urbanum* (right panel) sampled from 39 populations in Berwickshire.

